# Functional design of bacterial superoxide:quinone oxidoreductase

**DOI:** 10.1101/2021.12.23.473985

**Authors:** Abbas Abou-Hamdan, Roman Mahler, Philipp Grossenbacher, Olivier Biner, Dan Sjöstrand, Martin Lochner, Martin Högbom, Christoph von Ballmoos

**Affiliations:** Department of Chemistry, Biochemistry and Pharmaceutical Sciences, University of Bern, 3012 Bern, Switzerland; Institute of Biochemistry and Molecular Medicine, University of Bern, 3012 Bern, Switzerland; Department of Plant and Microbial Biology, University of Zürich, 8008 Zürich, Switzerland; Stockholm center for Biomembrane Research, Department of Biochemistry and Biophysics, Stockholm University, 10691 Stockholm, Sweden

## Abstract

The superoxide anion - molecular oxygen reduced by a single electron - is produced in large amounts by enzymatic and adventitious reactions and can perform a range of cellular functions, including bacterial warfare and iron uptake, signalling and host immune response in eukaryotes. However, it also serves as precursor for more deleterious species such as the hydroxyl anion or peroxynitrite and therefore, cellular defense mechanisms for superoxide neutralization have evolved. In addition to the soluble proteins superoxide dismutase and superoxide reductase, recently the membrane embedded diheme cytochrome *b*_*561*_(CybB) from *E. coli* has been proposed to act as a superoxide:quinone oxidoreductase. Here, we confirm superoxide and cellular ubiquinones or menaquinones as natural substrates and show that quinone binding to the enzyme accelerates the reaction with superoxide. The reactivity of the substrates is in accordance with the here determined midpoint potential of the two *b* hemes (+48 and -23 mV / NHE). Our data suggest that the enzyme can work near the diffusion limit in the forward direction and can also catalyse the reverse reaction efficiently under physiological conditions. The data is discussed in context of described cytochrome *b*_561_ proteins and potential physiological roles of CybB.

## Introduction

In all aerobic organisms, oxygen plays a vital role in the conversion of reducing equivalents into the universal cellular energy currency ATP. During the last step of respiration, terminal oxidases catalyze the highly exergonic reduction of molecular oxygen to water. This reaction is coupled to the formation of a transmembrane electrochemical proton gradient (or proton motive force, *pmf*), which is harnessed by the ATP synthase to recycle ATP from ADP and inorganic phosphate. In contrast to this well controlled reaction, adventitious single electron transfer to molecular oxygen results in formation of reactive oxygen species (ROS), representing incompletely reduced oxygen variants such as superoxide (O_2_·^-^), hydrogen peroxide (H_2_O_2_) and hydroxyl radical (OH^.-^). Depending on their concentration, ROS can either be toxic or function as signalling molecules, playing important roles in normal physiological and pathophysiological processes^1^. High and sustained concentrations of ROS are known to have deleterious effects on cellular health by oxidizing the three major biomolecule classes DNA, proteins, and lipids^2^. As molecular oxygen preferably partitions into the lipophilic environment of the membrane, the electron transfer reactions of the membrane embedded respiratory chain enzymes are a prominent source of ROS, e.g. at the flavin sites of complex I^3^ and II^4^ and the ubiquinol oxidation center of complex III^5^. Single electron transfer to molecular oxygen yields superoxide, which shows relatively mild deleterious cellular effects, but has been shown to directly damage the metal centers of enzymes^6^. In eukaryotes, superoxide is actively produced by the enzyme NAPDH oxidase as a second messenger, but also to serve as a first line of defense against bacterial infections^7-9^. In prokaryotes, extracellular superoxide production has been observed in marine bacteria to support metal uptake^10^, and as well as in *Enterococcus faecalis, E. coli* and *Vibrio cholerae*^*11-13*^. The main defense mechanism is production of superoxide dismutase, first described in 1969, which greatly accelerates the disproportionation of two superoxide molecules to one molecule of hydrogen peroxide and one molecule of oxygen^14,15^. Some twenty years later, a second enzyme, superoxide reductase, was described in anaerobic bacteria to react specifically with superoxide, forming hydrogen peroxide using reduced rubredoxin as electron donor^16^.

Yet another 20 years later, in 2018, our groups described a third, membrane-embedded enzyme, encoded by the *Escherichia coli* gene *cybB* with the gene product known as cytochrome *b*_561_ (CybB). In contrast to the two other enzymes, CybB catalyzes the oxidation of superoxide to oxygen and transfers the extracted electron to the oxidized quinone pool in the membrane^17^ (Fig. 1A). Therefore, CybB is a superoxide:ubiquinone oxidoreductase or, in short, superoxide oxidase (SOO). The crystal structure and functional data suggested that in the forward reaction, superoxide binds to a patch of positively charged amino acid residues at the periplasmic surface. This triggers a transmembrane electron transfer reaction via two *b*-type hemes, and the reduction of ubiquinone bound near the cytoplasmic surface of the protein (Fig. 1A). The two hemes show a close edge-to-edge distance of 11 Å allowing a fast electron transfer^18,19^. When CybB was mixed with reduced ubiquinol, the reverse reaction, i.e., reduction of CybB hemes and production of superoxide, was observed. The direction of catalysis under physiological conditions and the metabolic role of CybB are unknown. However, *mRNA* levels of *cybB* were found to be more than 10-times higher in the exponential than in the stationary phase whilst being unaffected by the presence of oxygen^17^.

**Figure 1:**
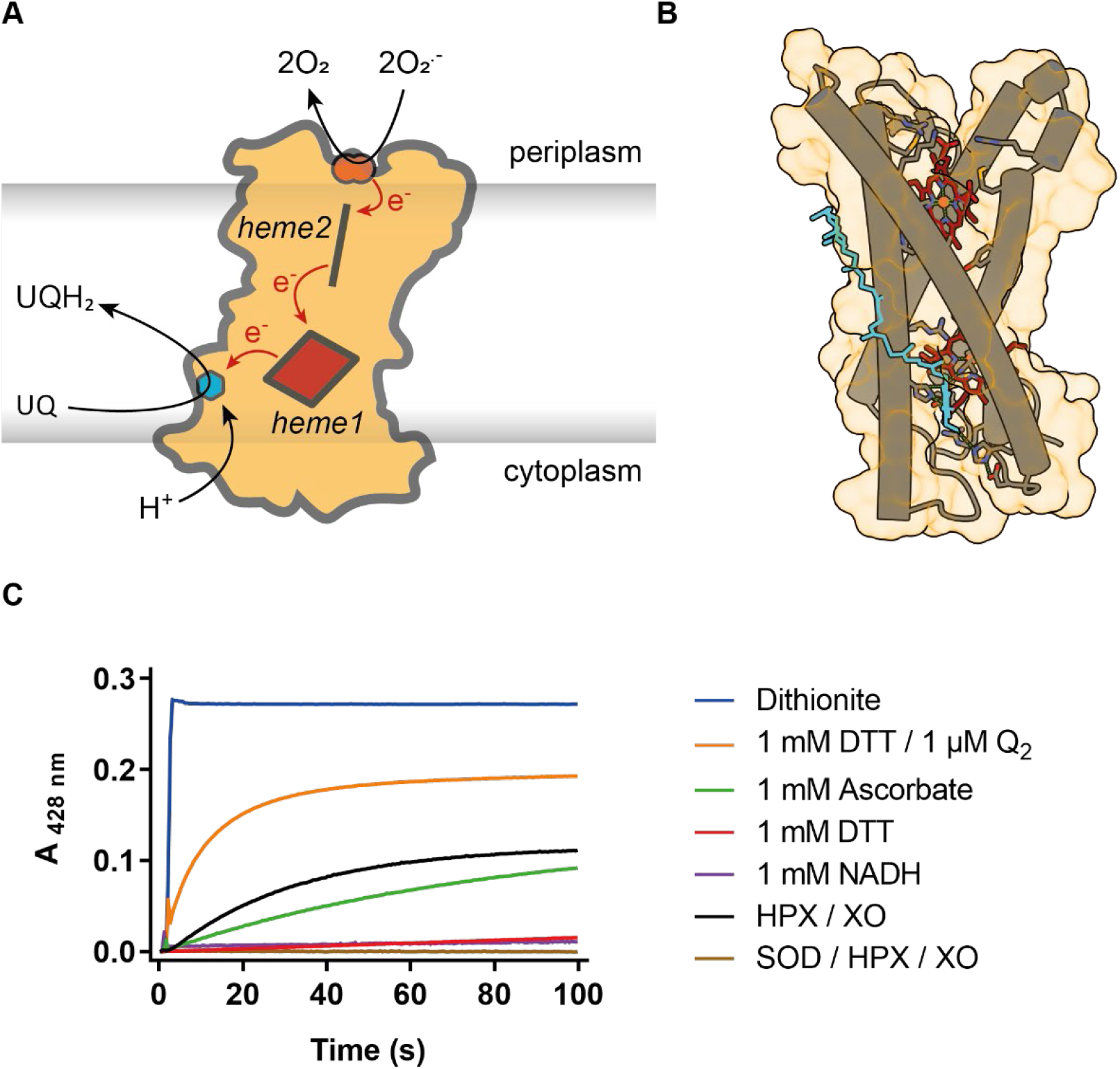
CybB structure and the corresponding substrates. **A)** Proposed reaction mechanism of CybB. Upon oxidation of superoxide anion at the heme 2, electrons are transferred through the two hemes to reduce the membrane quinone pool with protons being sequestrated from the cytoplasm **B)** Distorted helical arrangement of CybB produced by ECSF Chimera X. The 4 alpha helices are shown in grey, heme 2 and heme1 in dark red and ubiquinone UQ8 in cyan. Visualization of ubiquinone Q_8_ (PubChem CID 5283546) as suggested by molecular docking to the crystal structure (pdb 5OC0) using AutoDock Vina^25^. **C)** Kinetic traces monitoring the CybB reduction at 428 nm upon addition of different substrates: 1 mM sodium dithionite (blue), 1 mM dithiothreitol (DTT) (red), 1 mM DTT / 1 uM ubiquinone 2 (Q_2_) (orange), 0.1 mM Hypoxanthine / 7.6 mU Xanthine oxidase (HPX / XO) (black), 1 mM Ascorbate (green) and 1 mM NADH (purple). Measurements were done in 100 mM sodium phosphate pH 8, 0.1 mM DTPA, 0.05 % DDM. When xanthine oxidase is used, 0.1 mM of hypoxanthine was added to the previous buffer. 8.8 µg ml^-1^ of SOD was present before addition of substrates (brown).

Here, we thoroughly characterize detergent-solubilised superoxide:quinone oxidoreductase from *E. coli* using a range of biochemical and biophysical techniques. We provide evidence that superoxide and cellular quinone are indeed its natural substrate and measure the pH dependence of the reverse reaction. The data are then interpreted in connection to the heme mid-point potential (also determined in this study) in the absence and presence of quinone. Using a coupled enzymatic assay, we determine that the apparent affinity constant *K*_m_for quinones is in the submicromolar range.

## Materials and methods

### Expression of CybB

Expression of CybB was done as published previously^17^. Briefly, the plasmid p*cybB*-His encoding for the *E. coli cybB* gene with an 8xHis-tag fusion at its C-terminus was used to transform chemically competent *E. coli* BL21(DE3) pLysS cells. Transformed bacteria were grown on LB-agar plates containing 30 µg ml^-1^ kanamycin overnight at 37 °C. One colony was picked to inoculate a culture in 100ml LB media containing kanamycin (30 µg ml^-1^). Large culture growth was conducted at 37 °C using a LeX48 system (Epiphyte 3, Canada) where conventional shaking was replaced by continuous air bubbling. At a culture absorbance (OD_600_) between 0.9 and 1.1, expression of CybB was induced by the addition of 0.2 mM IPTG. Cells were harvested 4 hours after induction by centrifugation at 7500 x g and the pellet was resuspended in buffer (50 mM Hepes pH 7.5, 200 mM NaCl and 7.5 % glycerol).

### Membrane preparation and protein purification

To prepare membranes, the resuspended pellet was mixed with 1 mM PMSF, 2 mM MgCl_2_, some crystals of PEFA-bloc (Biomol, Germany), lysozyme and DNase I (Merck, Germany). The mixture was stirred for 30 min at RT and the homogenized cells were broken at least twice at 2000 Bar utilizing a Maximator High Pressure Homogenizer HPL6 (Maximator AG, Switzerland). Unbroken cells were removed by centrifugation at 7500 x g for 15 min and the supernatant transferred to a Beckman Ti45 ultracentrifuge rotor and centrifuged at 40000 rpm for 45 min. The membrane pellet was resuspended in the resuspension buffer and diluted up to 10 mg ml^-1^. Diluted membranes were solubilized by the addition of 1 % of OGNG (or 1 % DDM) (Anatrace) in the presence of 1 mM PMSF and some crystals of PEFA-Bloc for 1 h at 4 °C. Insoluble material was cleared by ultracentrifugation for 45 min at 40000 rpm in a Beckman Ti70 rotor. The resulting supernatant was loaded onto the nickel-charged Profinity IMAC Resin (Bio-Rad, USA) and washed with 15 column volumes (CV) of wash buffer (10 mM Hepes pH 7.5, 200 mM NaCl, 7.5 % glycerol and 0.1 % OGNG) and 15 CV of wash buffer 2 (10 mM Hepes pH 7.5, 200 mM NaCl, 7.5 % glycerol, 0.1 % OGNG and 5 mM Histidine). CybB was eluted with 10CV of elution buffer (10 mM Hepes pH 7.5, 200 mM NaCl, 7.5 % glycerol, 0.1 % OGNG containing either 200 mM histidine, 100 mM EDTA or 200 mM Imidazole) and concentrated up to 0.5 ml using Amicon Ultra-15 centrifugal filters with MWCO of 50 kDa (Merck, Germany). The sample was then applied to a size exclusion chromatography on a Superdex 200 increase 10 /300GL column equilibrated with wash buffer. Fractions containing CybB were pooled together, concentrated with a MWCO 50 kDa concentrator, and flash-frozen in liquid nitrogen.

### Reduction of CybB

Reduction of CybB was monitored using a Cary 60 UV-Vis Spectrophotometer (Agilent Technologies, USA) by measuring heme absorbance changes at either 428 nm or 561 nm upon addition of various substrates in aerobic conditions. All measurement were done in 100 mM sodium phosphate pH 8, 0.1 mM diethylenetriaminepentaacetic acid (DTPA) and 0.05 % DDM in the presence or absence of 0.1 mM hypoxanthine (HPX) depending on whether superoxide is produced by xanthine oxidase (XO) or not. In those measurements, 0.25 to 1 µM of CybB was used and the reaction started by addition of superoxide (generated enzymatically by using 0.01 U xanthine oxidase from bovine milk (Sigma, USA) that catalyse the oxidation of hypoxanthine to xanthine and / or xanthine to uric acid and produces superoxide as by-product, chemically by dissolving 7.1 mg KO_2_ in 1 ml of anhydrous DMSO in the presence of 80mg of 18-crown-6 ether to stabilize produced superoxide under steam of N_2_^20^ or upon using of 1 mM NADH / 10 µM PMS^21^) or other reductants such as 1 mM dithiothreitol (DTT) and 1 mM ascorbate. For control measurements, 8.8 µg ml^-1^ of superoxide dismutase (Sigma) was used to disproportionate superoxide anion in the presence of 20 µg ml^-1^ of catalase (Sigma). Full reduction of CybB was achieved upon addition of sodium dithionite and used hence to calculate the percentage of relative heme reduction. Ubiquinone analogues were either synthetized (Q_1_, Q_2_ and decylubiquinone) as described in the supplementary material or purchased from Sigma. Ubiquinone analogues (Q_1_, Q_2_ and decylubiquinone) and naphthalene analogues (1,4 naphthoquinone and menadione, a soluble form of menaquinone) were reduced by incubation of 50 mM quinone solution with 1 M DTT with respect of V_DTT_ / V_Q_ ratio of 4 to ensure a complete reduction of quinone or pre-reduced with sodium borohydride NaBH_4_ (as with Q_2_). Reduced quinones were hereafter added to reduce CybB and the percentage of relative quinol heme reduction was calculated as mentioned previously.

For the pH dependency measurements, we used the same protocol as before and measured either superoxide or Q_1_H_2_ induced heme reduction in the presence of different buffered solutions. In the first set of experiment, we used 100 mM sodium phosphate (pH 7 or 8) or 100 mM CHES and 100 mM sodium phosphate dibasic (pH 9 or 10) containing 20 mM KCl, 200 mM NaCl, 0.1 mM DPTA, 0.05 % DDM. In the second set, we used 100 mM sodium phosphate (for pH ranging from 6 to 8.5) or 20 mM Tris-HCl (for pH ranging from 9 to 10) containing each 20 mM KCl, 200mM NaCl, 0.1 mM DPTA, 0.05 % DDM. In both experiments, the percentage of relative heme reduction compared to a full dithionite reduction of CybB was calculated. Anaerobic measurements were performed after removal of dioxygen using a combined system of argon blowing in the presence of 53 µg ml^-1^ catalase, 26 µg ml^-1^ glucose oxidase and 5 mM glucose mixture (all purchased from Sigma).

### Oxidation of reduced CybB

Oxidation of reduced CybB was monitored under anaerobic condition. Anaerobicity was achieved as described above (section: *Reduction of CybB*). 15 min after argon blowing, 0.25 µM of CybB was fully pre-reduced using a tiny amount of dithionite (6.6 µM in 1.5 ml of 100 mM sodium phosphate pH 8, 20 mM KCl, 200 mM NaCl, 0.05 % DDM) before addition of 10 µM of diverse quinone (ubiquinone and naphthalene analogue) to oxidize it. A full oxidation of reduced CybB was achieved by adding 10 µM of potassium ferricyanide and used to calculate the percentage of relative heme oxidation.

### Superoxide assay and *K*_*m*_ determination

We used the previously established WST-1 (Water soluble Tetrazolium dye) assay^22^ and adapted it^17^ to determine either the apparent affinity constant for ubiquinone of CybB or to measure superoxide production by the enzyme. In both assays, formazan formation upon reduction of WST-1 by superoxide was monitored at 438 nm in assay buffer (100 mM sodium phosphate pH 8, 0.1 mM DTPA, 0.1 mM HPX, and 0.05 % DDM) containing 50 µM WST-1, 20 µg ml^-1^ catalase and 60 nM *bo3* oxidase. Measurements were done either individually in a cuvette using Cary 60 UV-Vis spectrophotometer or in a 96-well plate in SpectraMax ABS Plus Microplate Reader (Molecular Devices, USA). Ubiquinone Q_1_ or Q_2_ were added to the previous mixture to measure the overall activity of CybB. The apparent Michaelis Menten constant (*K*_*m*_) for quinone was calculated using a constant amount of CybB (50 nM) while varying the quinone concentration between 0 and 100 µM. To measure superoxide production and determine the apparent constant affinity for quinol, the reaction was initiated by addition of 50 nM of CybB to the assay buffer containing different concentration of quinol (0 - 100 µM) in the absence of *bo*_*3*_ oxidase and catalase.

The second order rate (catalytic efficiency) of CybB was calculated based on the known value for SOD (2 × 10^9^ M^-1^ s^-1^)^23^. Briefly, the concentration for both enzymes SOD and CybB was titrated using the former assay to reduce the formazan formation rate by 50 %. The determined concentrations were used to calculate the catalytic efficiency of CybB.

### Potentiometric redox titrations

Redox titrations were performed anaerobically at room temperature in a Spectro-electrochemical cell (Prosense, Netherlands), by following heme absorption changes upon a gradual addition of dithionite solution as electron source, instead of electrochemistry as in^24^ using a UV-Vis spectrophotometer.

Throughout the measurement, anaerobicity was maintained by a constant and continuous flow of argon. Prior to measuring, 10 μM CybB, in the absence or presence of 20 µM Q_1_, and 5 μM mediators were incubated together in 100 mM sodium phosphate pH 7, 0.1 mM DTPA, 0.1 mM HPX, and 0.05 % DDM and flushed with a repetitive and alternative cycles of N_2_ / vacuum to ensure complete removal of oxygen. The following mediators were used: benzyl viologen (−350 mV), sodium anthraquinone 2-sulfonate (−225 mV), 2-hydroxy 1,4-naphtoquinone (−145 mV), resorufin (−51 mV), methylene blue (+11 mV), phenazine ethosulfate (+55 mV), phenazine methosulfate (+80 mV), 2,6-Dichlorophenol-indophenol (+217 mV) and tetramethyl-p-phenylendediamine (+276 mV). Redox potentials were measured using an Ag / AgCl 3 M KCl reference electrode associated to a platinum counter electrode (Pt) (Prosense, Netherlands). The redox potential was noted after equilibration, and an optical spectrum was recorded between 400-600 nm. The measured absorbance of heme reduction at 561nm was plotted against the measured redox potentials and curves were fitted using the Nernst equation with n=2 due to two successive one electron reductions corresponding to the reduction of both hemes. All midpoint potentials referenced here were adjusted by the addition of +210 mV relative to the normal hydrogen electrode (vs NHE) at 25 °C.

## Results

### Enzyme production and purification

The gene sequence of *E. coli cybB*, equipped with a C-terminal octahistidine tag was cloned into a pET-28b vector under control of the *lac* promoter. As a host strain, BL21(DE3) pLysS was used. The protein is well expressed yielding cells and inverted membranes with a distinct pink color. Here, we further optimized the purification of CybB based on our published protocol^17^. The membrane fraction was solubilized with either 1 % OGNG or 1 % DDM and purified via Ni-IDA chromatography. The protein was eluted using either imidazole, histidine or EDTA, concentrated and assessed using size-exclusion chromatography (Superdex 200 Increase, 10 /300GL, 500 μl injected), using either 0.1 % OGNG or 0.1 % DDM. The results displayed in supplementary Figure S1 show a major impact of both the detergent and the Ni-IDA eluant used. While both detergents extracted the protein similarly from the membrane, only extractions with OGNG produced a monomeric peak (∼14.7 ml), as judged from the size exclusion chromatograms (Fig. S1 A and E). In addition to this peak (Fig S1A), two further discrete elution peaks were observed, indicating aggregate forms of the protein. If the protein was eluted with either 100 mM EDTA or 200 mM histidine (Fig. S1 A and E) instead of imidazole a homogenous preparation with a single monomeric peak was obtained. In the following, 1 % OGNG and 200 mM histidine in 0.1 % OGNG was used for solubilization and elution, respectively.

### Spectroscopic properties and reaction with cellular reductants

The crystal structure of CybB shows a highly distorted 4-helix bundle with the *b*-hemes fully coordinated by histidines (Fig. 1B). The protein is almost entirely in contact with the lipid bilayer with no peripheral domains. Heme 2 is partly exposed towards the periplasm by a narrow cavity that contains a variety of positively charged residues that are proposed to guide the negatively charged superoxide anion in proximity of the heme to allow electron transfer. Heme 1 is located towards the cytoplasm and buried in the membrane with no polar access route, supporting the idea of a binding site for a hydrophobic compound such as ubiquinone. The two hemes are rapidly reduced by dithionite and the reduced spectrum shows a characteristic Soret peak at 428 nm as well as α and β peaks at 530 and 561 nm as described earlier. However, all other attempts to fully reduce the enzyme with cellular reductants or ubiquinols failed. Lundgren et al^17^. found that detergent solubilized CybB is rapidly, but only partly reduced (∼50 %) by the DTT-reduced short-chain ubiquinone analogue Q_1_. Partial reduction by superoxide was also only observed when using xanthine oxidase as superoxide producing system. No or only very slow reduction is observed with reducing agents such as ascorbate, DTT and NADH (Fig. 1C). We therefore aimed to better understand the reduction behaviour of detergent solubilized CybB in a series of kinetic and steady state experiments.

### *Reaction of CybB with superoxide produced by KO*_*2*_ and NADH / PMS

Superoxide in aqueous solution has a half-life that is strongly pH dependent, as the spontaneous disproportionation to hydrogen peroxide and oxygen occurs via the protonated O_2_H form. Consequently, superoxide is more stable at higher pH values, i.e., t1/2∼1 s at 10 μM and pH 8^26^. If superoxide was indeed the reducing agent, the reaction of superoxide with CybB should be more complete at higher pH values. To test this, KO_2_ dissolved in anhydrous DMSO stabilized with 18-crown-6 ether was used to produce a stable superoxide stock solution. From this solution, a small amount was added to an aqueous solution containing detergent solubilized CybB (1 µM) and the reduction of CybB was followed spectrophotometrically (mixing time ∼2 s) at 428 nm at different pH values (Fig. 2A). Both the rate and the level of reduction increased with increasing pH correlating with a prolonged presence of superoxide at higher pH values (Fig. 2 A, B). Essentially full reduction was observed at pH values ≥10. If the reaction at pH 10 was supplemented with SOD (8.8 µg ml^-1^), reduction of CybB was suppressed, indicating the superoxide specificity of the system. A further system that has been described to produce superoxide is NADH mixed with the electron mediator phenazine methosulfate (PMS)^21^. Indeed, when this mixture was incubated with detergent solubilized CybB, rapid reduction of the hemes was observed and the reaction was completely suppressed in the presence of SOD (Fig. 2C).

**Figure 2:**
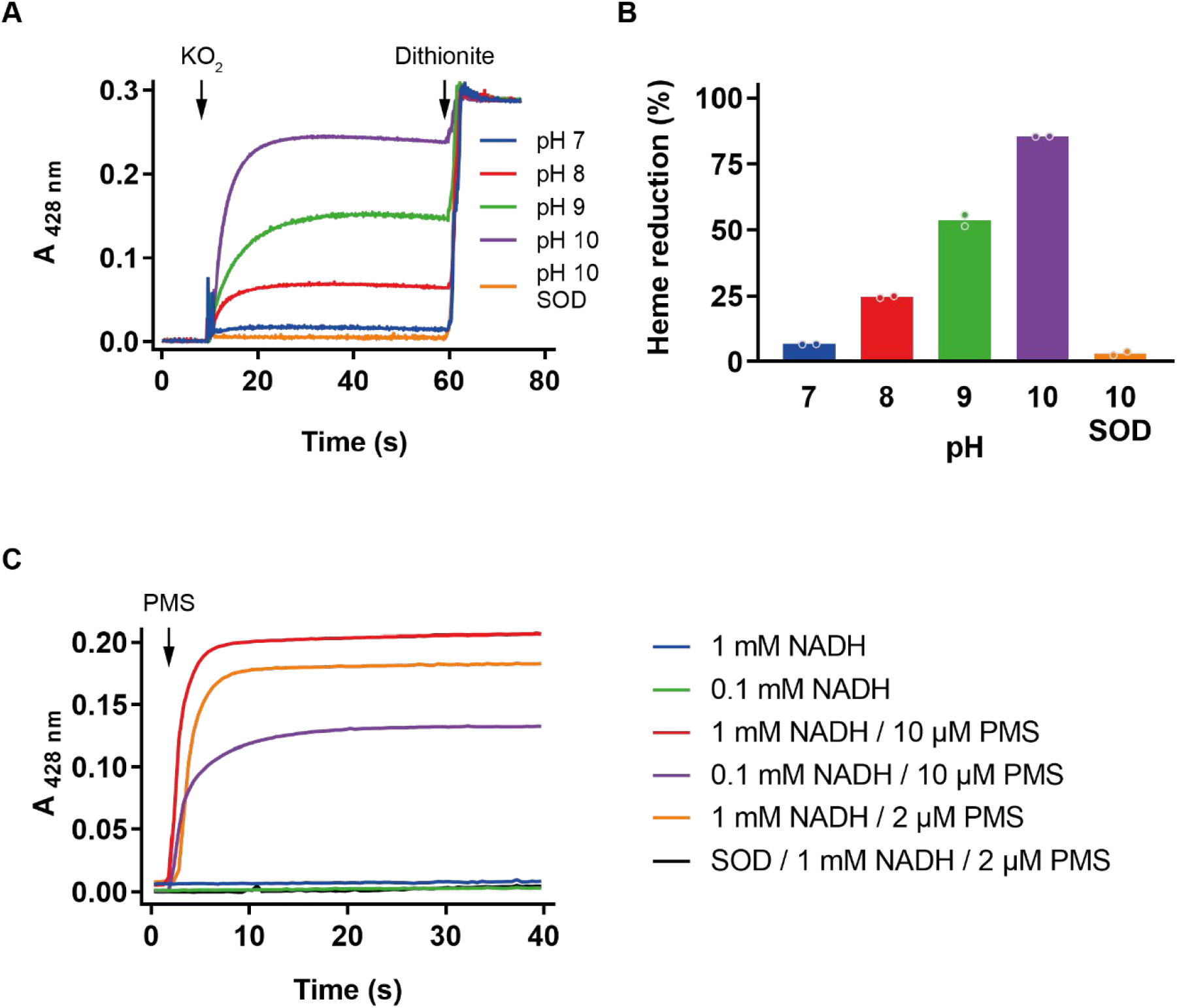
Superoxide CybB reduction. *A) Kinetic traces monitoring CybB reduction at 428 nm upon addition of 5 µM superoxide from KO*_*2*_ *solution at different pH values. Full reduction of CybB was achieved by addition of sodium dithionite. B) percentage of the relative level of reduction after 20 s incubation with KO*_*2*_ *(Data from Fig. 2A). C) Kinetic traces monitoring CybB reduction in presence of various amount of NADH and PMS at pH 8*. NADH and SOD (8.8 µg ml^-1^) for the controls were added from beginning, PMS was added after 2 s.

### Reaction of CybB with reduced ubiquinol

To investigate the reverse reaction, we mixed detergent solubilized CybB (1 μM) with ubiquinol QH_2_, prereduced with either NaBH_4_ or *in situ* by 1 mM DTT 1 min prior to addition of the enzyme. CybB reduction by DTT alone is very slow and can be ignored (Fig. 3A). As depicted in Figure 3A, rapid reduction is observed if solubilized CybB is mixed with 1 mM DTT and 10 μM ubiquinone Q_2_ (purple trace). If 1 μM Q_2_is used, the rate is slightly lower, but the same level of reduction is reached. At 0.1 μM Q_2_, the reaction was slowed down >10 times, and the maximum reduction was not reached within the measuring period. Next, we varied the pH of the reaction mixture, keeping, a more water-soluble analogue of Q_2,_ DTT / Q_1_, constant at 1 mM / 10 μM. As depicted in Fig. 3B, reduction levels of CybB were higher at alkaline pH values, while essentially no heme reduction was observed at pH 6 or below. We had previously observed that upon the reaction with quinol, the enzyme consumes oxygen and produces superoxide, as detected with WST-1^17^ (see also Fig. 6E). Consequently, under aerobic conditions, the enzyme is turning over, and the observed reduction level reflects steady-state catalysis. Therefore, the experiment was repeated under anaerobic conditions and while a similar trend was found, reduction levels were increased at all pH values compared to aerobic conditions (Fig. 3B). The difference between aerobic and anaerobic reduction was most pronounced at pH values <7.5. A plausible explanation for this behaviour is that the level of reduction correlates with the midpoint potential of the two reactants, the *b* hemes of CybB and ubiquinone. The latter is strongly pH dependent (∼-60 mV per pH unit), as two protons participate in the reaction^27^. A higher pH value thus corresponds to a lower midpoint potential, thus increasing the reducing power, while the midpoint potentials of protein embedded hemes are expected to be less sensitive to pH changes^28^.

**Figure 3:**
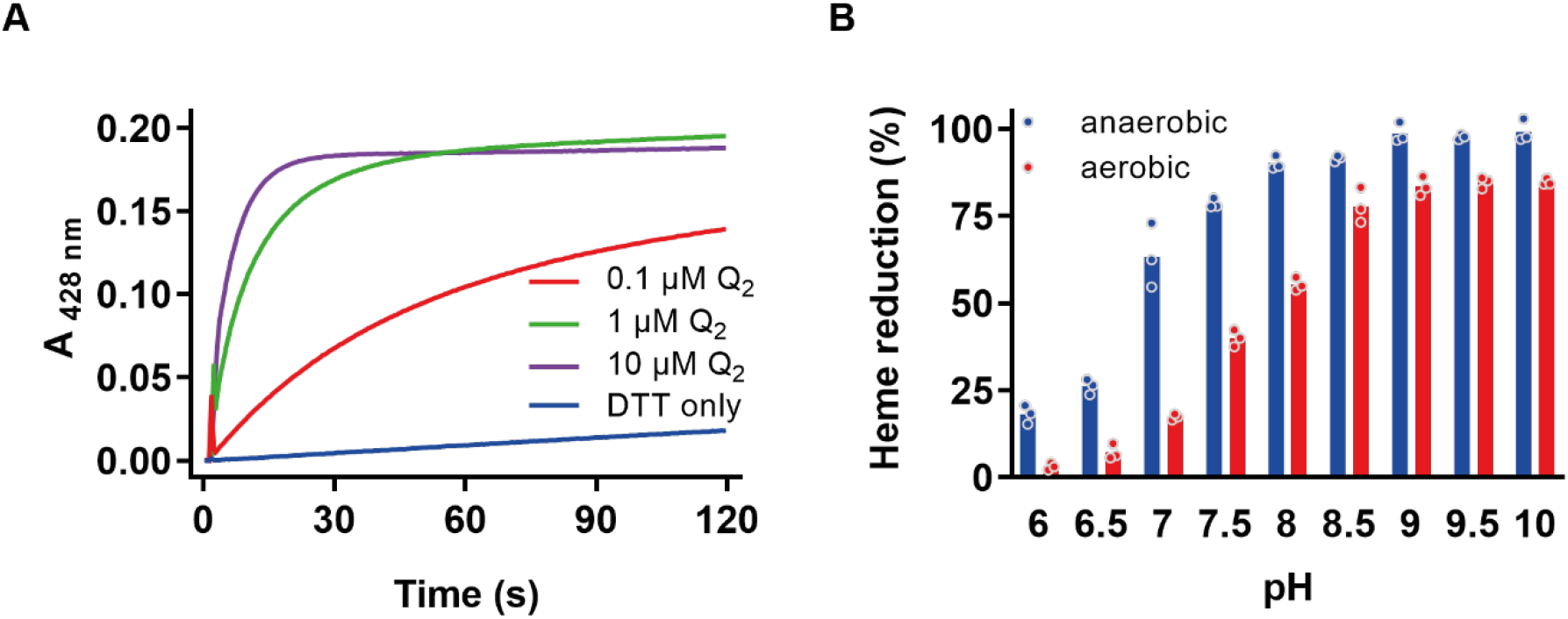
Quinol CybB reduction. **A)** Kinetic traces monitoring CybB reduction at 428 nm upon addition of DTT alone or premixed with different amounts of ubiquinone Q_2_ at pH 8. **B)** percentage of heme reduction measured at 561 nm after 20 s incubation of CybB with 1 mM DTT / 10 μM Q_1_ at different pH values under aerobic and anaerobic conditions.

### Redox titration of CybB

In 1986^29^, the midpoint potential of the two hemes was determined with the purified enzyme and a single value of ∼20 mV was obtained at pH 7.4. However, it is unlikely that both hemes have an identical mid-point potential. If the forward reaction was favourable, a lower midpoint potential is expected for the periplasmic heme 2 compared to the cytoplasmic heme 1 to support electron transfer from superoxide via heme 2 and heme 1 to ubiquinone. We therefore performed a redox titration with dithionite under anaerobic conditions in the presence of a set of electron mediators and followed the potential with a voltmeter and the reduction state spectrophotometrically at 561 nm. Several titrations were done in the absence and presence of ubiquinone Q_1,_ and the obtained traces, depicted in Figure 4A, were fitted for two hemes. The two waves have similar amplitudes, each of them reflecting 50 % of the relative reduction signal as expected for two hemes with similar absorption properties. In the absence of ubiquinone, midpoint potentials of -23 ± 12 mV and + 48.5 ± 12.5 mV NHE were determined, close to the reported values, and we assigned them to heme 2 and heme 1, respectively (Figure 1B). If ubiquinone Q_1_is present, the two potentials are shifted, one from – 23 mV to – 8 mV and the other from 48.5 to 100 mV. Importantly, the difference in midpoint potentials is increased (108 mV), suggesting that binding of ubiquinone thermodynamically facilitates electron transfer from heme 2 to heme 1 (Fig. 4B). In addition, the increase of the potential at heme 2 facilitates reduction by superoxide, while the potential at heme 1 (100 mV) is still fully compatible with quinone reduction. From these measurements, it can be speculated that binding of quinone to the oxidized enzyme could influence the reaction of the enzyme with superoxide. We therefore incubated CybB either with stoichiometric amounts or a 10-fold excess of quinone, before superoxide production was initiated by the addition of xanthine oxidase. As depicted in Figure 4C, the observed reaction rate of the preincubated enzyme with superoxide increased dramatically. The reduction kinetics in presence of ubiquinone was biphasic, with a fast first phase and a slower second phase of reduction, which likely can be contributed to the reduction of CybB by ubiquinol (formed by either the adventitious reaction of superoxide with quinone or by CybB itself). To test this hypothesis, ubiquinol *bo*_3_oxidase was added to keep the ubiquinone pool oxidised and prevent reduction by ubiquinol. The addition of *bo*_3_ oxidase resulted in a monophasic reduction of CybB with the first phase only, indicating that reduction occurred via superoxide only.

**Figure 4:**
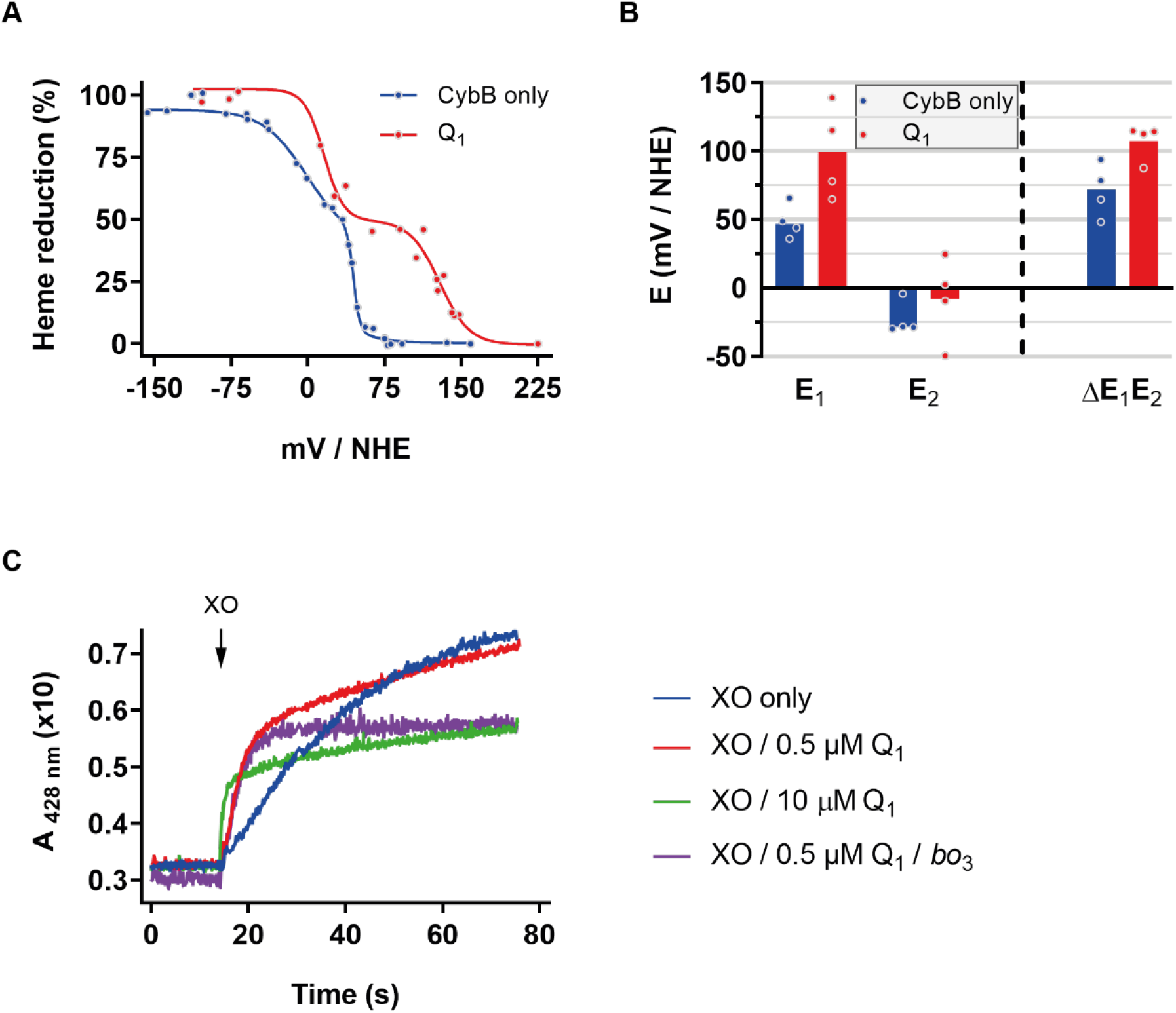
Heme redox potentials. **A)** Midpoint redox potential titrations curves of 10 µM CybB in the absence (black) and in the presence (red) of 20 μM Q_1_. Redox potentials were performed anaerobically in 100 mM sodium phosphate pH 7, 0.1 mM DTPA, 0.1 mM HPX and 0.05 % DDM and redox potentials are expressed in mV against NHE. **B)** Energy diagram of potentials calculated from fits of Fig. 4A. Shown are the values for the low (E_2_) and high potential heme (E_1_) and their energy difference(ΔE_1_E_2_) in the presence and absence of quinone. **C)** Kinetic traces showing reduction of 0.5 µM of CybB by superoxide anion produced by xanthine oxidase in the absence and in presence of an equimolar (0.5 µM) or an excess amount (10 µM) of ubiquinone Q_1_ and 60 nM of bo_3_ oxidase. Measurement was done in 100 mM sodium phosphate pH 8, 0.1 mM DTPA, 0.1 mM HPX and 0.05 % DDM.

### Both ubiquinone and menaquinone are substrates of CybB

In contrast to the reverse reaction, the forward reaction is more complicated to monitor due to the short half-life of superoxide and its hard-to-control production. We have therefore simplified the system and pre-reduced CybB under anaerobic conditions with stoichiometric amounts of dithionite, until ∼90-95 % of the enzyme is reduced to avoid presence of unreacted dithionite.

Addition of ubiquinone 10µ M Q_1_ showed a rapid oxidation of CybB, indicating that ubiquinone is a substrate (Fig. 5A, red trace). A similar behavior was found with other short-chain ubiquinones such as Q_2_ and decylubiquinone. Next, we tested the two menaquinone analogues menadione and 1,4-naphtoquinone. Reaction with 1,4-naphtoquinone proceeded also quickly, but enzyme oxidation was less complete, while the reaction with menadione was >10 slower and less enzyme was oxidized. A summary of the different experiments is found in Figure 5B, showing that reaction with ubiquinones reached almost complete CybB oxidation, in reference to the oxidation by ferricyanide. Addition of air saturated water also led to oxidation of CybB, indicating that the reduced enzyme can deliver electrons to oxygen in the absence of quinones.

**Figure 5:**
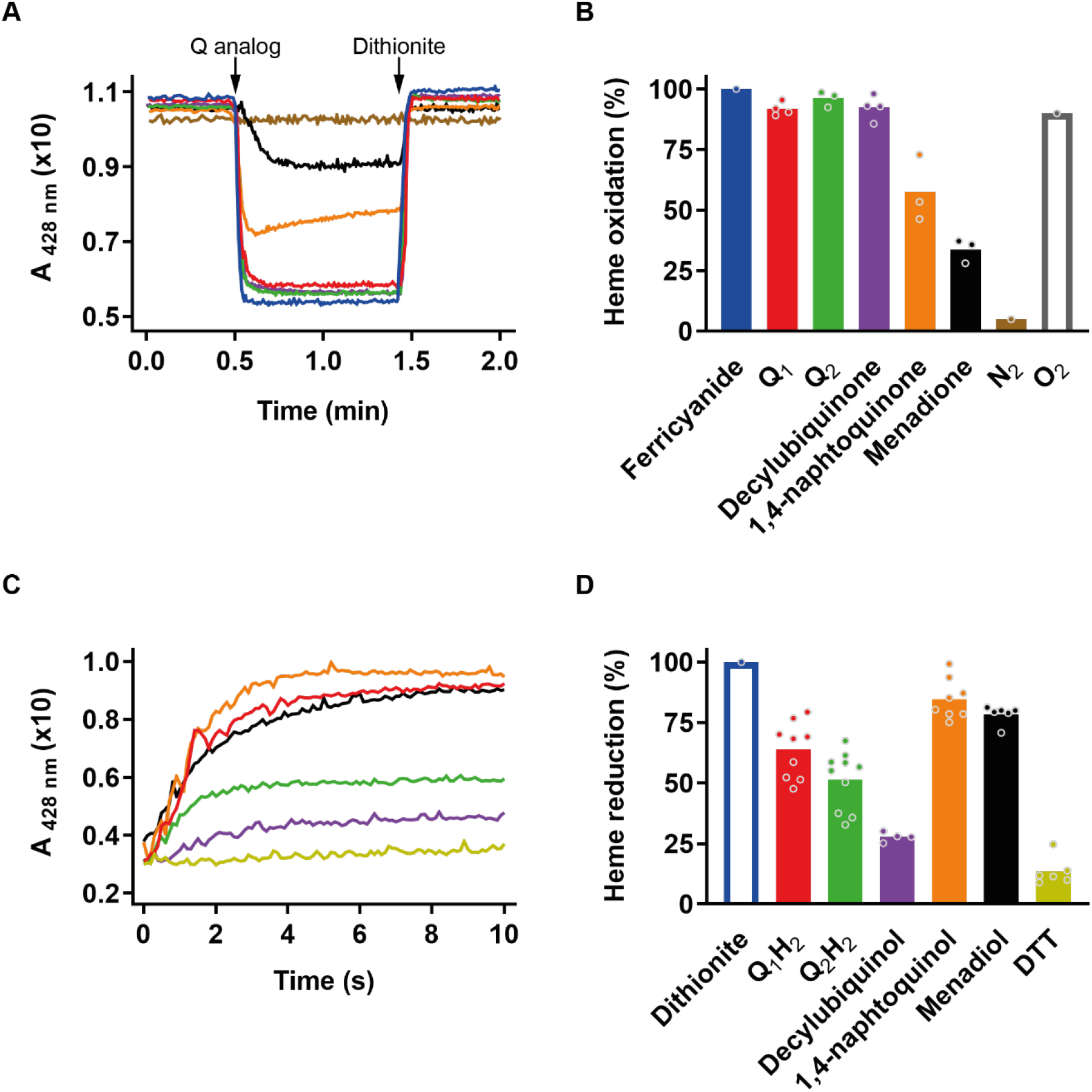
Ubiquinone and menaquinone as substrates for CybB. **A)** Kinetic traces of CybB oxidation as observed at 428 nm. CybB was reduced, under anaerobic conditions, with small amounts of dithionite (6.6 µM) until >90 % reduction was obtained, after which 10 µM of different quinones was added. After the reaction was complete, CybB was fully reduced by addition of 3.3 mM dithionite. **B)** Summary of the percentage of reduced heme oxidation, including the data from addition of air-saturated water (not included in Fig. 1A). **C)** Kinetic traces of CybB reduction, under aerobic conditions, as observed at 428 nm. Different quinols were added at 10 μM concentration. **D)** Summary of the results obtained from Fig. 5C. (Note that the reference sample (Dithionite) has no trace in 5C). The bars in figures 5B and 5D have the same color as the respective oxidized (5A) or reduced (5C) quinone traces.

Next, we compared the ability of the reduced form of these different quinones to reduce CybB under aerobic conditions, i.e., the reverse reaction. When using ubiquinol Q_1_H_2_at pH 8, ∼60 % of reduction of CybB was observed as discussed above (Fig. 5C). Again, the observed kinetics were very similar to Q_2_H_2_and decylubiquinol, but both quinols showed a lower level of reduction. In contrast to the forward reaction, incubation with the reduced menaquinol analogues led to almost complete reduction of the enzyme (Fig. 5 C, D). Hence, the observed data correlate well with our hypothesis that the lower midpoint potential of menaquinones (compared to ubiquinones), leading to a more complete reduction of CybB.

### Measurement of steady state activity of the forward reaction and determination of quinone affinity

Having the two most likely substrates identified, we aimed to develop a turnover assay that reports the actual protein function, converting superoxide to reduced quinol. However, it is not trivial to directly measure quinol formation (see discussion). We therefore adapted a superoxide dismutase assay using WST-1, a water-soluble indicator for superoxide, which after reaction with superoxide is converted into a stable formazan that strongly absorbs at 438 nm^17^. As a constant superoxide producing system, xanthine oxidase and hypoxanthine was employed, which in phosphate buffer has been reported to produce ∼50 % superoxide and 50 % hydrogen peroxide^30,31^. Although neither CybB nor WST-1 has been shown to react with hydrogen peroxide, catalase is included in the assay buffer to avoid accumulation of hydrogen peroxide. Consequently, the linear increase observed at 438 nm represents the continuous conversion of WST-1 by xanthine oxidase produced superoxide into the stable formazan (Fig. 6B, blue trace). Winterbourn and others have previously used this assay to estimate SOD content of tissue extracts, as any present SOD competes with WST-1 for superoxide leading to a decreased formazan formation^22^. We adapted this idea and added the required components for CybB to compete with WST-1 for superoxide. If CybB (50 nM) and Q_2_ (10 µM) were present (orange trace on Fig. 6), the initial signal was a bit lower, but quickly regained the initial slope. We interpreted this result as competition between CybB and WST-1 for superoxide that stops after enough ubiquinone has been converted to ubiquinol. We therefore added purified *E. coli bo*_3_quinol oxidase to the mixture to regenerate the quinone pool, which led to a continuous suppression of formazan production (red trace on Fig. 6). Importantly, if *bo*_3_and Q_2_ were present in the absence of CybB, no suppression was observed (green trace on Fig. 6). This indicates that the spontaneous reaction of superoxide with ubiquinone to semiubiquinone - accompanied by the disproportionation to quinone and quinol (that can be consumed by the *bo*_3_ oxidase) - is not sufficient to explain the observed suppression. The overall scheme of the assay with all components is depicted in Figure 6A.

**Figure 6:**
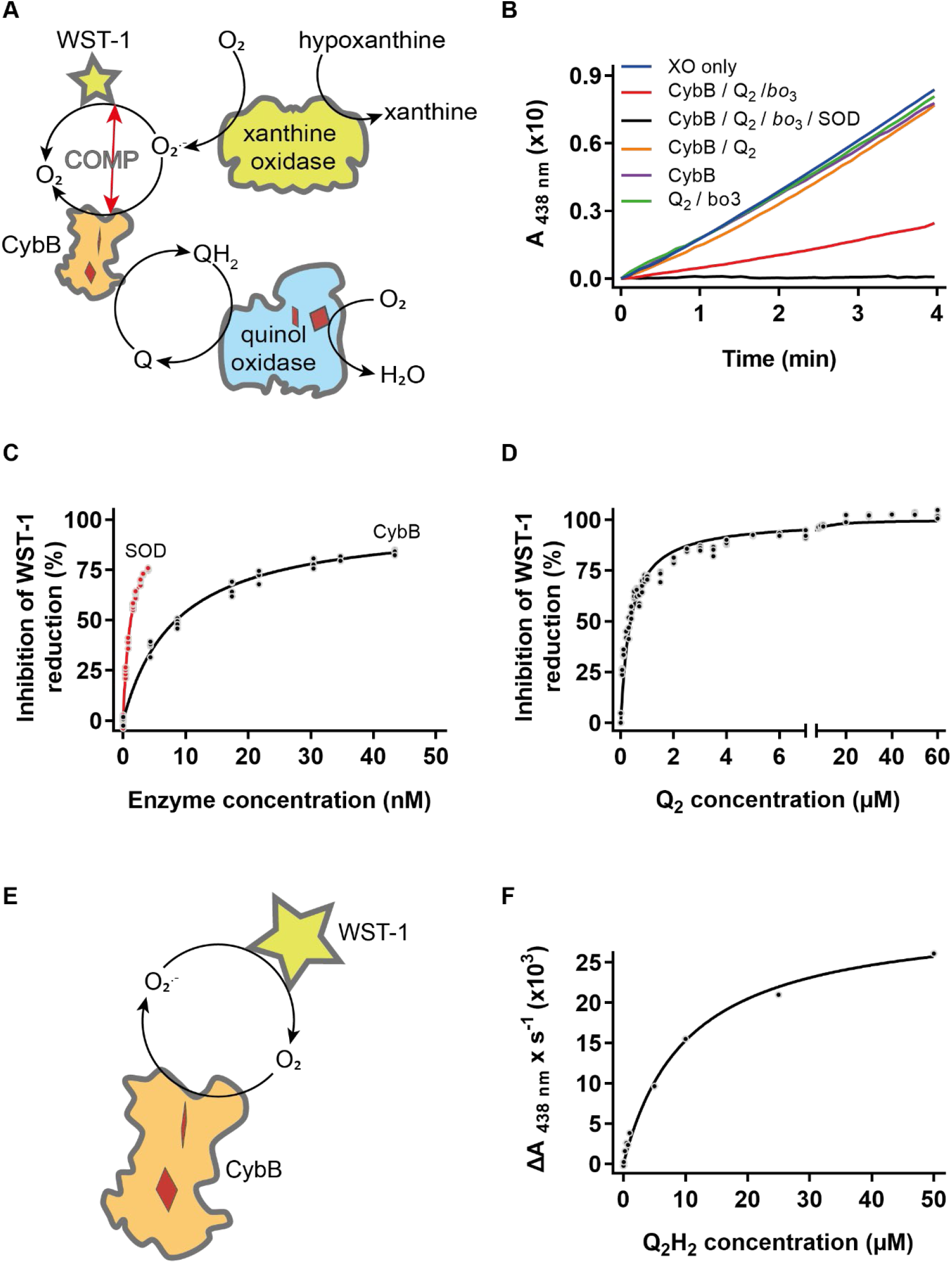
Superoxide assay and apparent K_m_ determination. **A)** Cartoon depicting the coupled assay to follow competition (COMP) between CybB and WST-1 for superoxide. **B)** Kinetic traces monitoring formazan formation at 438 nm. The different conditions are indicated in the legend. **C)** Titration of different amounts of purified CybB and purified SOD under otherwise identical experimental conditions (0.01 U xanthine oxidase, 10 μM Q_2_, 60 nM bo_3_ oxidase). Relative decrease in formazan rate formation (inhibition of WST-1 reduction %) upon addition of either SOD (red trace) or CybB (black trace). **D)** Determination of the apparent K_m_ of ubiquinone Q_2_ in the forward reaction. Relative decrease in formazan rate formation upon addition of different amounts of Q_2_. **E)** Cartoon depicting the assay scheme to determine reverse activity, i.e., formation of superoxide by quinol. **F)** Determination of apparent K_m_ of ubiquinol Q_2_H_2_ in the reverse reaction. Initial slopes of formazan formation (ΔA _438nm_ x s^-1^ (×10^3^)) upon addition of different amounts of prereduced ubiquinol Q_2_H_2_ (as depicted in Supplementary Fig. S3).

We used this approach to estimate both the second order rate constant of the reaction and the apparent *K*_*m*_ for ubiquinone. First, in the presence of all components including 10 μM Q_2_ and *bo*_3_ oxidase, different amounts of SOD and CybB were added and the decrease in formazan formation was followed. As depicted in Figure 6C, about 10-times more CybB was required to obtain similar quenching values. Taking the previously determined second order rate constant of 2 × 10^9^ M^-1^ s^-1^ for SOD, a rate constant in the range of 1 to 3 × 10^8^ M^-1^ s^-1^ can be estimated for CybB (see table 1). To measure the apparent *K*_*m*_ of the reaction for ubiquinone, all components except ubiquinone were mixed, and a steady formazan formation was monitored for 30 s, before different amount of Q_2_ were added from stock solutions, leading to decreased levels of formazan formation (Figure 6D). Using Michaelis-Menten kinetics, an apparent *K*_*m*_ of 390 ± 40 nM was obtained. We determined very similar *K*_*m*_values by either subsequent addition of increasing amounts of quinone as well as with different quinone concentrations measured in parallel in a 96-well plate.

**Table 1:**
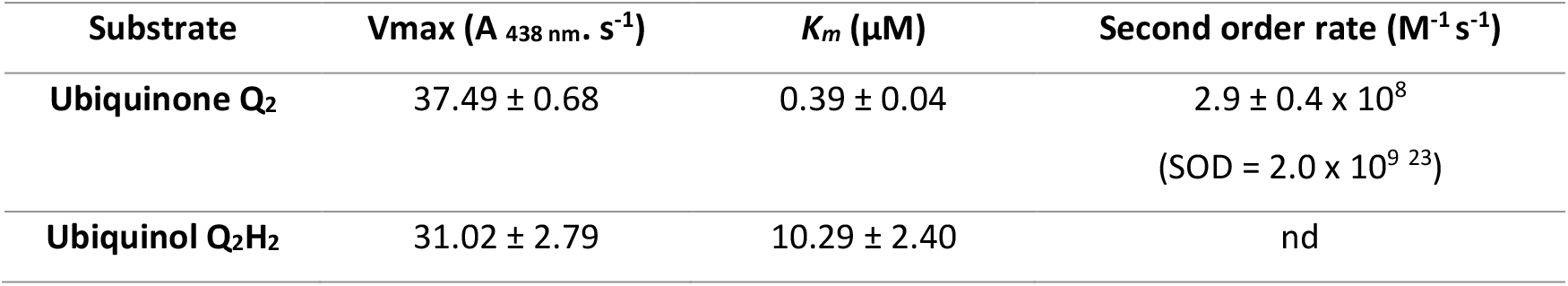
Summary of kinetics parameters as determined using WST-1 assay.

| Substrate | Vmax ($A_{438\text{ nm}} \cdot s^{-1}$) | $K_m$ ($\mu M$) | Second order rate ($M^{-1} s^{-1}$) |
| --- | --- | --- | --- |
| Ubiquinone $Q_2$ | $37.49 \pm 0.68$ | $0.39 \pm 0.04$ | $2.9 \pm 0.4 \times 10^8$<br>(SOD = $2.0 \times 10^9$<sup>23</sup>) |
| Ubiquinol $Q_2H_2$ | $31.02 \pm 2.79$ | $10.29 \pm 2.40$ | nd |

Finally, we also determined an apparent *K*_*m*_ for the reverse reaction. To this end, superoxide formation by CybB was monitored following formazan formation from WST-1 (Fig. 6E and supplementary Fig. S3) after addition of NaBH_4_-prereduced ubiquinol. As depicted in Figure 6F, if initial rates were plotted against quinol concentrations, an apparent *K*_*m*_ of ∼10 µM was obtained, thus being ∼25 higher than for the forward reaction (see table 1).

## Discussion

### Architecture of a superoxide: quinone oxidoreductase

The enzyme described here, CybB or cytochrome b_561_ from *E. coli* was proposed to act as a membrane bound superoxide:ubiquinone oxidoreductase^17^. It is only the third enzyme described, after SOD and SOR, to specifically react with superoxide. However, in contrast to the two other enzymes that contain a catalytic dimetal center, CybB is a hemoprotein containing two *b* hemes. Superoxide produced in biological systems is either the product of an enzymatic or adventitious reaction of an electron with molecular oxygen. It has a standard midpoint potential of -160 mV, which however is expected to be much higher under biological conditions as the O_2_/O_2_·^-^ ratio is expected to be very large in the cell (e.g., a ratio of O_2_/O_2_·^-^ the potential increases to ∼+42 mV). Superoxide spontaneously disproportionates into oxygen and hydrogen peroxide, which are more stable products. In principle, superoxide can act as a mild reductant, however, it is better known as an oxidant oxidizing lipids or enzyme metal centers^6^. Superoxide reacts with many organic molecules in solution, including ubiquinone. The product, ubisemiquinone, has been shown to disproportionate into ubiquinone and ubiquinol^32^. However, this spontaneous reaction is not expected to occur in biological systems, as superoxide is considered membrane impermeable due its negative charge, while ubiquinones are tightly embedded in the membrane. Superoxide production can also be envisioned to happen at the membrane surface as oxygen preferentially partitions into the membrane, and quinone binding sites of respiratory enzymes have been described as point of superoxide formation^3,33-35^. In a similar scenario, water-soluble ascorbate and membrane-bound ubiquinone are unable to react in cells, even though they have been shown to rapidly react in solution^36^. The proposed design of a membrane embedded superoxide:quinone oxidoreductase therefore makes sense if electron transfer between the two otherwise separated substrates should happen. The orientation of CybB in the membrane predicts that reduction occurs from the periplasm via the hemes to the quinone binding site located on the cytoplasmic side. The idea of a quinone binding site is consistent with the glycerol found in the crystal structure and docking studies^17^. The binding of superoxide from the periplasmic side is supported by an easily accessible heme from the protein surface surrounded by positively charged amino acids. Such a positive patch would attract, and guide, negatively charged superoxide anion radical O_2_·^-^, creating an entry path for superoxide.

Similar positive patches have also been found in the human Cu,Zn-SOD^37^, *Alvinella pompejana* Cu,Zn-SOD^38^ or the superoxide reductase (SOR) of *Desulfoarculus baarsii* in complex with Fe(CN)_6_^30,40^. However, the strongest indication for a specific intercatio006E of superoxide with CybB is the recent structure of the transmembrane domain of the NADPH oxidase (Nox5) from *Cylindrospermum stagnale*^7,41^. The protein consists of a transmembrane domain formed by six helices, with an overall similar fold as CybB, harboring two *b* hemes in close distance. The protein protrudes into the cytoplasm with an NADPH binding domain containing FAD as cofactor. This topology allows the enzyme to use cellular NADPH to produce superoxide and hydrogen peroxide on the extracellular side to act as chemical weapon in cellular warfare to combat bacterial infections. Notably, the two proteins, Nox5 and CybB, show a very similar electrostatic environment around the periplasmic heme, which is surrounded by positively charged amino acids creating an easy access route for negatively charged species in the periplasmic compartment (see Fig. 7). In particular, it has been proposed that Arg256 present in this cavity could electrostatically promote the catalytic production of superoxide^7^. The overall reaction of Nox5, oxidation of NADPH and reduction of oxygen to superoxide is thus very similar to the reverse reaction of CybB, ubiquinol oxidation and superoxide formation, and the structural arrangement of the two enzymes, especially with respect to the exposed heme and the positive patch, is also strikingly similar.

**Figure 7:**
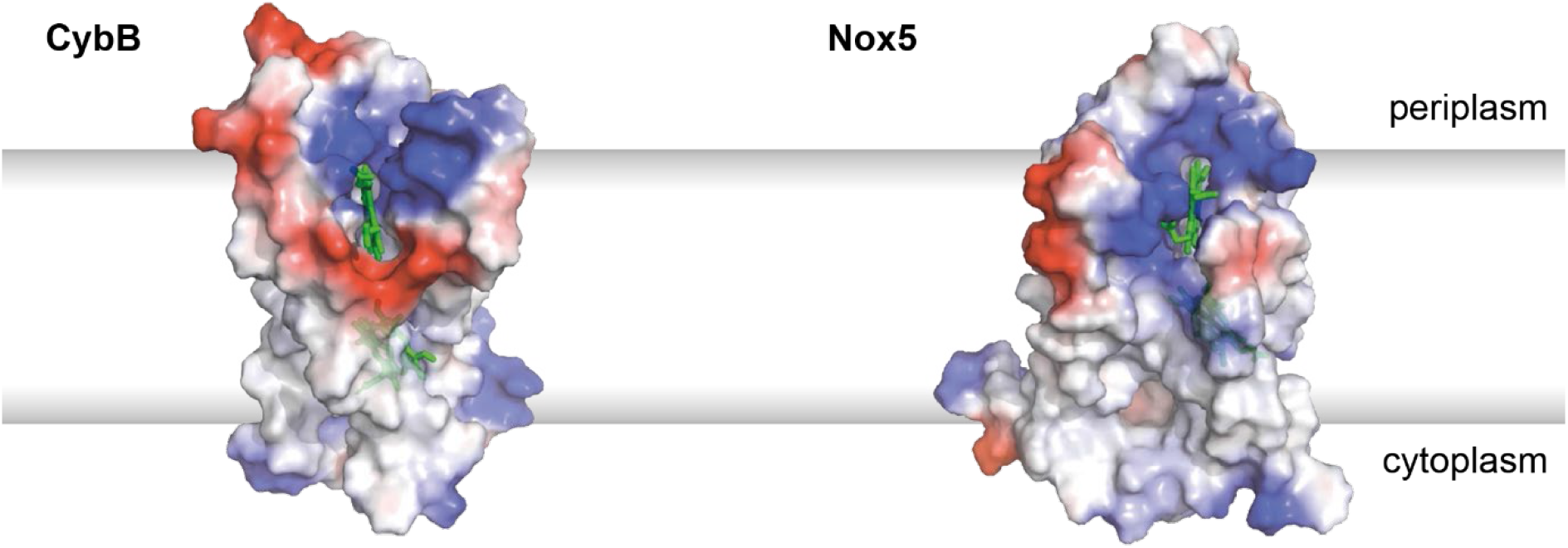
Comparison of the electrostatic potential map. *The charges distribution around the exposed edge of heme group (green) of CybB (pdb 5OC0) and Nox5 (pdb 5O0T) show a similar positive surface potential (blue) that is presumed to funnel the superoxide anion to the heme edge. Images were produced in Pymol*.

### Reaction of CybB with its substrates

A short description of enzyme purification is included in the results section for a reason. Even though the purified CybB was very pure on SDS-PAGE gel immediately after purification via Ni-IDA resin and the reduced dithionite spectra appeared identical to those after gel filtration, functional characterization revealed different kinetics and levels of heme reduction in different enzyme preparations when reduced with DTT / Q1 or superoxide. We therefore performed the described experiments by varying detergent and IDA elution methods, until we found conditions that yielded protein with reproducible properties. Possible reasons for the inconsistent behaviour in the other preparations are different levels of delipidation and protein aggregation. It can be expected that a small protein with such a distorted structure relies on interactions with its annular layer of lipids, and this requires further investigation in the future.

With the optimized enzyme preparation in our hands, the experiments presented here provide strong evidence that superoxide and both ubiquinone and menaquinone are natural substrates of CybB. This quinone substrate diversity is not surprising, as all terminal *E. coli* oxidases (*bo*_3_, *bdI* and *bdII*) have been shown to accept either UQ_8_, DMQ_8_ or MQ_8_^42^, as well as *E. coli* the nitrate reductase (NarGHI) during nitrate respiration^43-45^. We show that CybB is readily reduced by short chain ubiquinols in a strongly pH dependent manner. The standard midpoint potential of ubiquinone / ubiquinol is ∼70 mV at pH 7^46^, but decreases by – 60 mV per pH unit, rendering it a stronger reductant at higher pH values in accordance with our data. We also find that the observed steady state reduction is increased under anaerobic conditions, in agreement that in the absence of oxygen the enzyme cannot turnover leading to an accumulation of a reduced intermediate. We show that menaquinols, that have a lower midpoint potential (∼-100 mV), reduce the enzyme more completely than ubiquinol. The opposite picture is observed, if the prereduced enzyme is mixed with the oxidized form of the different quinone analogues under anaerobic conditions. While incubation with oxygen and ubiquinone led to a completely oxidation comparable to ferricyanide, incubation with the lower potential menaquinone led only to a partial oxidation of the enzyme, corroborating that the observed level is dependent on the redox potential of the substrate. In contrast to ubiquinone, the redox potential of the two heme groups are not expected to be heavily pH dependent, as no proton takes place in the redox reaction.

The specific reaction of CybB with superoxide is more complex to analyse because of difficulties to produce sufficient concentrations due to the instability of superoxide. At pH 8, the half-life of 10 μM superoxide is approximately 1s ^26^. Here, we have used three well-established methods to produce superoxide in aqueous solutions, whilst superoxide specificity was tested by signal suppression after the addition of SOD^22,47,48^. First, superoxide was produced enzymatically by xanthine oxidase, which is routinely used to investigate SODs. However, it is nearly impossible to quantify the superoxide directly, as the percentage of superoxide produced per hypoxanthine molecule varies strongly and depends on the buffer conditions and on the enzyme preparation itself^31^. The activity of commercially available xanthine oxidase must thus be analysed for every purchased batch. Second, we used inorganic potassium dioxide mixed with a crown-ether that produces a stable superoxide solution in organic solvents, e.g., DMSO or acetonitrile and finally we used NADH and phenazine methosulfate (PMS), which produces superoxide in aqueous solution^21^. All three systems rapidly reduced CybB and, in all cases, addition of SOD suppressed CybB reduction. To exclude unspecific reaction of superoxide with other heme-containing enzymes, we incubated purified *bo*_3_ oxidase with the xanthine oxidase / hypoxanthine and found essentially no heme *b* reduction over the time course of CybB reduction (Supplementary Fig. S2). In the case of KO_2_, we analysed the pH dependency of the reaction and found that CybB can be fully reduced at higher pH values. Since the redox potential of the oxygen / superoxide couple is not proton dependent, we attribute this observation to the increased stability of the superoxide anion (and hence a higher substrate concentration). It is important to point out that all tested low redox potential substrates such as NADH (−340 mV), DTT (−330 mV), glutathione (−240 mV), lactate (−160 mV) or hypoxanthine (−400 mV) were unable to reduce the enzyme at pH 8. Earlier reports that NADH and lactate reduce CybB in membrane vesicles are likely due to adventitious superoxide production during respiration by NADH and lactate dehydrogenases, respectively^49^. A notable exception is ascorbic acid (+60 mV), described as natural substrate for many other cytochrome *b*_561_ proteins, which reduced CybB of *E. coli* slowly and in a pH dependent fashion (no reduction below pH 7), likely due to its decreased potential at higher pH values. Interestingly, in cytochrome *b*_561_ of secretory vesicles that regenerate internal ascorbate from semidehydroascorbate via transmembrane electron transfer, the reduction of the enzyme is not pH sensitive between pH 6 and 8, indicating that a different mode of action with ascorbate is utilised in this enzyme compared to CybB^50^.

### Midpoint potential of CybB hemes

We determined the midpoint potential of the two hemes by titrating dithionite to an anaerobic solution of CybB in the presence of electron mediators. While Murakami et al^29^ found a single potential at +20 mV in a similar experiment, we were able to fit the titration to two waves with ∼50 % signal amplitude each, indicating that the two hemes have different potentials, i.e. -23 mV and +48.5 mV. These values are close to the ones reported by Murakami and well within the range found for other diheme *b* enzymes, i.e. heme *b* in *E. coli bo*_3_ oxidase and nitrate reductase have been shown to be +50 mV^51^ and +20 mV^52^, respectively. For diheme proteins of the cytochrome *b*_561_ family, typically slightly higher potentials have been found. In cytochrome *b*_561_ from bovine adrenal chromaffin vesicles, the redox potentials of high and low potential hemes were determined to be +150 and +60 mV, respectively^53^. A value of +80 mV ± 30 mV has been reported for the human duodenal cytochrome *b*_561_ that catalyse transmembrane electron transfer from external ascorbate to intravascular Fe^3+ 54^. In plants, the reported midpoint redox potential cytochrome *b*_561_ proteins range between +80 to +190 mV for the high potential heme and between -20 to +60 mV for the low potential heme, in accordance with the potentials of the proposed substrate couple dehydroascorbate / ascorbate (+60 mV)^55^. In the case of CybB, the measured midpoint redox potentials of –23 mV and +48.5 mV are assigned to periplasmic side faced heme 2 and cytoplasmic faced heme 1, respectively (Figure 1A). Repetition of the potential measurements in the presence of 20 µM ubiquinone Q_1_primarily shifted the midpoint potential of heme 1 to higher values increasing the potential difference between the two hemes from +64 mV to +108 mV, supporting fast intramolecular electron transfer^56^. We thus hypothesize that binding of the substrate quinone close to heme 1 increases its potential, facilitating electron transfer from heme 2 to heme 1. We tested this hypothesis by comparing the reaction of CybB with superoxide in the presence and absence of quinone. In the presence of saturating quinone concentration the reaction was faster than the mixing time (t_1/2_<1 s). This effect is in accordance with the observation that binding of the semiquinone analogue inhibitor ‘HQNO’ decreased the potential of the low potential heme in succinate: menaquinone oxidoreductase from *Bacillus subtilis*^57^. Future experiments using EPR spectroscopy will help to measure and assign heme potentials reliably and pulse radiolysis studies will enable fast reaction kinetics of CybB with superoxide.

### Enzyme turnover measurements

The strongest evidence for the proposed reaction was obtained in a coupled enzyme assay that shows that CybB actively competes with WST-1 for superoxide and that this competition is dependent on the CybB and quinone concentrations, as well as on quinone regeneration by *bo*_3_ oxidase. This assay has been used and validated to detect SOD activity in tissues and purified fractions. Here, we confirm that only in the presence of all components, suppression of formazan formation was observed. If all components but CybB was present, WST-1 reduction by xanthine oxidase was unaffected. This is noteworthy, as it has been described that superoxide reacts with ubiquinone to form semiquinone, which can disproportion to ubiquinol and oxygen. Alternatively, semiquinone can react with oxygen to form superoxide again. However, if this reaction occured at a significant rate, formazan formation would already be decreased in the presence of quinone and *bo*_3_ oxidase (without CybB), which was not the case. Whether the added detergent affected this reaction, or the disproportion reaction was slow compared to the reaction of semiquinone with oxygen is unclear. To eliminate potential pitfalls, we have used the exact same conditions, including the presence of quinone and *bo*_3_ oxidase, and performed the experiment with purified SOD. We were surprised to find that only 10x more CybB than SOD was required to suppress WST-1 formation to a similar extent. This suggests that the second order rate constant of CybB is about 10 times lower than the one of SOD, yielding a still impressive value of ∼2.9 × 10^8^ M^-1^ s^-1^ in the presence of 10 μM ubiquinone Q_2_(see table 1). Such a rate makes CybB a true competitor for cellular superoxide even in the presence of SOD (see below).

This same assay was used to titrate quinone as a substrate, rendering a *K*_*m*_ of ∼390 nM, which is lower than what has been found for other quinone reducing enzymes in bacteria (typically in the micromolar range)^58-62^. Our data was analysed using Michaelis-Menten, which might be an oversimplification, as we do not know the “excess” of substrate superoxide over enzyme, which is further influenced by the spontaneous disproportion of superoxide. Obviously, such high turnover rates require a fast association of CybB with superoxide and a fast reduction of ubiquinone, and the rates of these have not been measured yet. In future experiments, pulse radiolysis will be used as an efficient mean to rapidly produce defined amounts of superoxide and investigate pre-steady-state kinetics on the enzyme as well as stopped-flow measurements to follow the oxidation of the enzyme by quinone. However, it might well be that superoxide binding to the quinone-bound enzyme is rate limiting, and subsequent electron transfer might be very fast, and that CybB intermediates with reduced hemes are thus not accumulating. The prerequisite of the quinone binding to CybB before the electron transfer from superoxide takes place might reflect a strategy to suppress reactive intermediates in the membrane. Mechanistically, it is important to remember that two molecules of superoxide must bind to fully reduce quinone to quinol and that protons must be taken up before reduced quinol can be released to the membrane and be used as a substrate for e.g., respiration. Future mutagenesis studies should allow us to identify the residues that are critical for these processes.

### Concluding remarks

The presented data strongly support our initial hypothesis, which was based on the crystal structure and a set of functional experiments^17^ that CybB of *E. coli* can work as a superoxide oxidase *in vitro*. While fast pre-steady state measurements like those performed for SOD and SOR are still outstanding, we find the present set of data corroborates our initials findings to be convincing. The measured redox potentials of the two hemes are lower than those of other cytochrome *b*_561_ proteins, suggesting a different substrate than ascorbate. In addition to the proposed forward reaction, also the reverse reaction (formation of superoxide from ubiquinol) is readily observed, and the similarity of the structural arrangement to the NADPH: superoxide oxidase is intriguing. The apparent *K*_m_ for quinone is 25 times lower in the forward direction, and for quinol in the reverse direction, making the forward reaction kinetically favourable.

To date, the physiological role of CybB is unknown, as simple growth experiments have not yielded a phenotype in a knockout strain. It seems unlikely, however that the forward reaction takes place under aerobic conditions, when the oxygen / superoxide ratio is high (>10^6^ in a laboratory culture). Under such conditions, the reverse reaction is more likely to occur. In this context, it is noteworthy that periplasmic superoxide formation has been described for *E. coli*^12^, and the reaction of menasemiquinone with oxygen has been suggested as a pathway for this. Based on our results with menaquinol, it is likely that CybB can catalyse such a reaction when the menaquinone pool is reduced during respiration, equipping *E. coli* with a “superoxide gun” to attack competitors. The idea is especially attractive since periplasmic SOD is mainly expressed during stationary phase, while CybB transcription is enhanced during the logarithmic phase^17^. Finally, the forward reaction can be envisioned to happen under microaerobic almost anaerobic conditions where the O_2_ / superoxide ratio is low; like the environment *E. coli* faces as a commensal bacterium of the human gut.

