## Supplementary Information for "Functional design of bacterial superoxide:quinone oxidoreductase"

**Affiliations:**

#### Supplementary Figures:

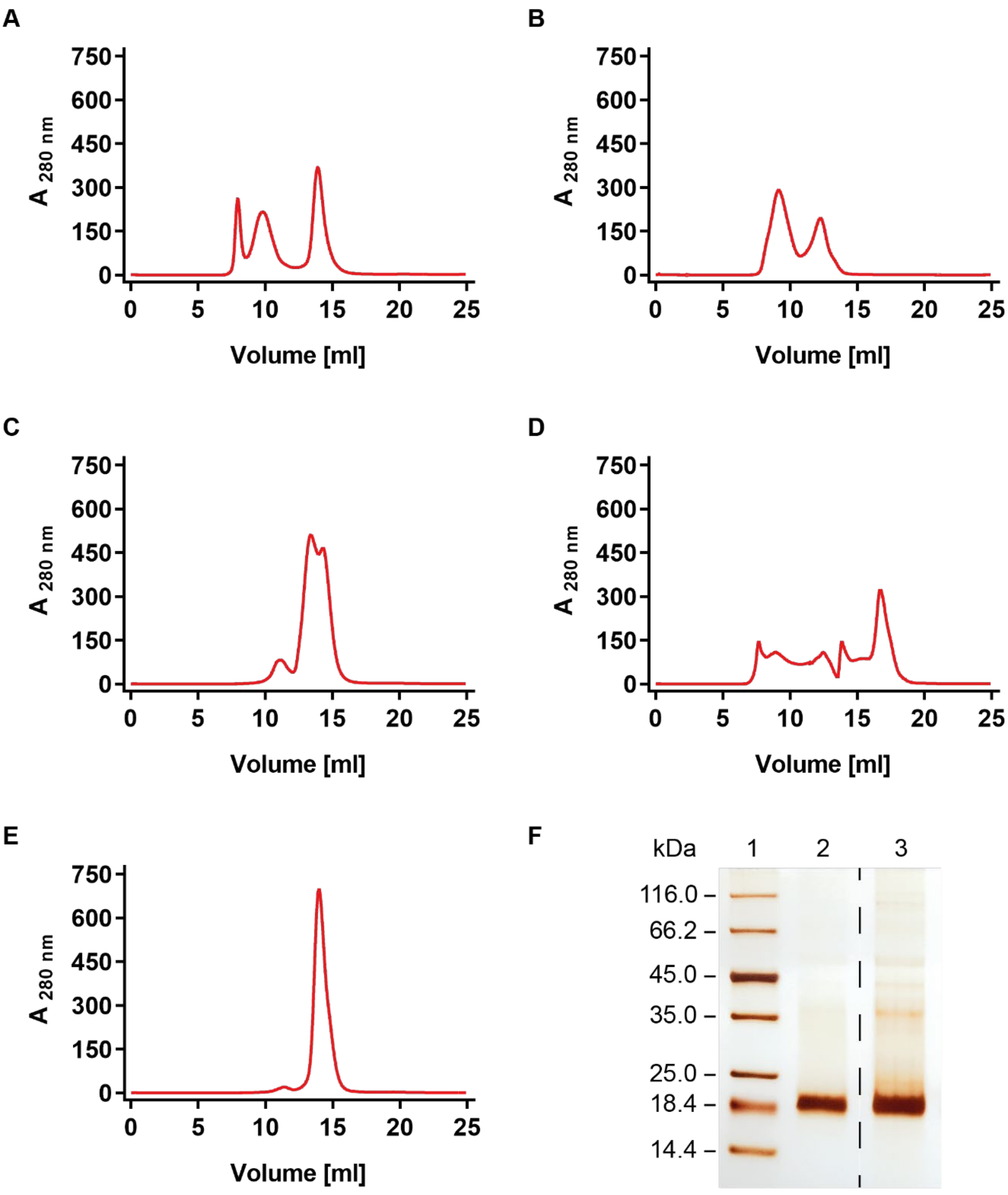

***Figure S1: Optimization of CybB purification. A-E)*** *Chromatograms showing CybB elution profile, upon injection on Superdex 200 increase 10/300GL, function of the nature of used detergent and eluant.* ***A)*** *Inverted Membrane (IM) were solubilized with 1 % OGNG and CybB was eluted with 100 mM EDTA.* ***B)*** *IM were solubilized with 1 % DDM and CybB was eluted with 100 mM EDTA.* ***C)*** *IM were solubilized with 1 % DDM and CybB was eluted with 200 mM histidine.* ***D)*** *IM were solubilized with 1 % OGNG and CybB was eluted with 200 mM histidine. OGNG was then exchanged to DDM during size exclusion chromatography.* ***E)*** *IM were solubilized with 1 % OGNG and CybB was eluted with 200 mM histidine.* ***F)*** *Comparison of the purity of CybB obtained after SEC after loading on 12 % SDS-PAGE gel as following: in line 1, Pierce$^{TM}$ unstained protein MW marker, in line 2, eluted protein from condition E and in line 3, eluted protein from condition C.*

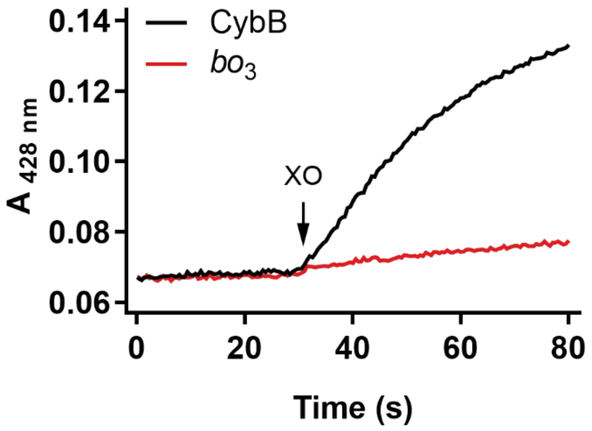

***Figure S2: reduction of heme group by superoxide.*** *Comparison of superoxide heme reduction between two heme containing enzymes (CybB and bo3 oxidase from E. coli). Absorbance changes was monitored at 428 nm and reaction was initiated by addition of 0.01 U of xanthine oxidase in 100 mM sodium phosphate pH 8, 0.1 mM DTPA, 0.1 mM HPX, 0.05 % DDM.*

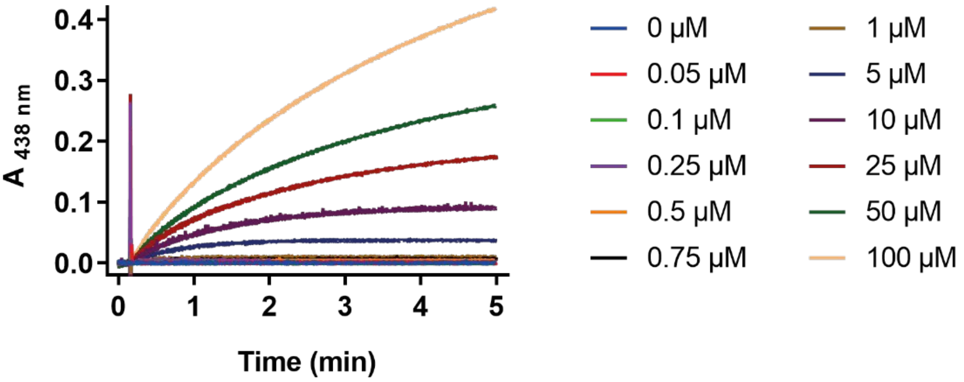

***Figure S3: Superoxide production by CybB.*** *Formazan formation was monitored at 438 nm upon reduction of WST-1 by superoxide anion produced by CybB in the presence of different amount of pre-reduced* $Q_2H_2$ *with sodium borohydride (0 - 100 µM). Reactions were initiated by addition of 50 nM CybB in 100 mM sodium phosphate pH 8, 0.1 mM DTPA, 0.1 mM HPX and 0.05 % DDM.*

### Synthesis of ubiquinone derivatives

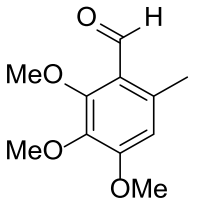

2,3,4-trimethoxy-6-methylbenzaldehyde
Chemical Formula: $C_{11}H_{14}O_4$
........ 8

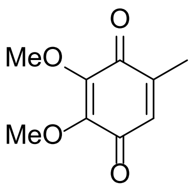

2,3-dimethoxy-5-methylcyclohexa-2,5-diene-1,4-dione
Chemical Formula: $C_9H_{10}O_4$
........ 10

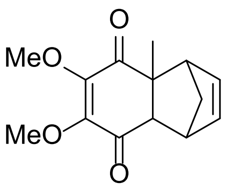

6,7-dimethoxy-4a-methyl-1,4,4a,8a-tetrahydro-1,4-methanonaphthalene-5,8-dione
Chemical Formula: $C_{14}H_{16}O_4$
........ 12

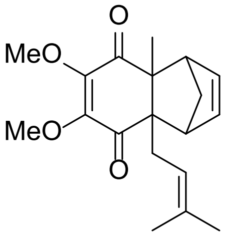

6,7-dimethoxy-4a-methyl-8a-(3-methylbut-2-en-1-yl)-1,4,4a,8a-tetrahydro-1,4-methanonaphthalene-5,8-dione
Chemical Formula: $C_{19}H_{24}O_4$
........ 14

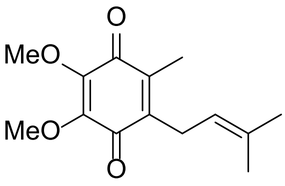

2,3-dimethoxy-5-methyl-6-(3-methylbut-2-en-1-yl)cyclohexa-2,5-diene-1,4-dione
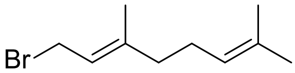
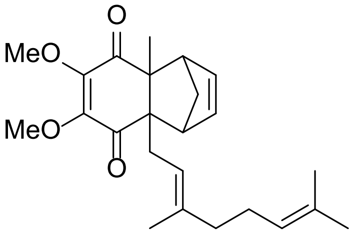
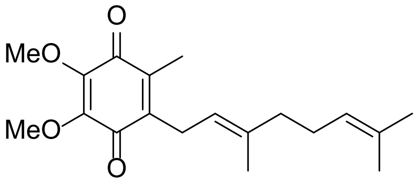
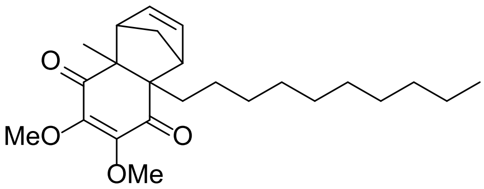
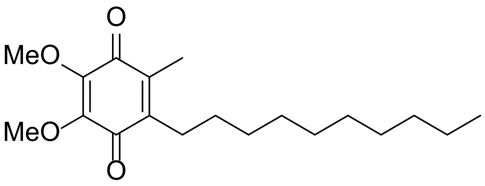

#### Methods

All reactions requiring anhydrous conditions were performed in heat-gun, oven or flame dried glassware under inert atmosphere ($N_2$, Ar). Ice baths were used to achieve 0 °C. For reactions at above room temperature heating blocks or PEG baths were used. Silica gel 60 Å (40–63 mm) from Sigma-Aldrich was used for dry loads. Flash column chromatography was performed on a Teledyne Isco CombiFlash® Rf+ with the corresponding RediSep® prepacked silica cartouches unless otherwise stated. Thin layer chromatography (TLC) was performed on Machery & Nagel Alugram® xtra SIL G/UV 254 visualization under UV light (254 nm) and/ or (366 nm) and/ or by dipping in anisaldehyde stain and subsequent heating. The synthetic protocols were modified from Lu and al,[1].

#### Chemicals

Commercial reagents and solvents (Acrôs Organics, Fluorochem, Grogg Chemie, Hänseler, Sigma-Aldrich) were used without further purification unless otherwise stated. Dry solvents for reactions were distilled and filtered over columns of dry neutral aluminium oxide under positive argon pressure. Solvents for extraction and flash chromatography were used without further purification.

#### Instrumentation

$^{1}H$ and $^{13}C$ NMR spectra were recorded on a Bruker AVANCE-300 or 400 spectrometer operating at 300 or 400 MHz for $^{1}H$ and 75 or 101 MHz for $^{13}C$ at room temperature unless otherwise stated. Chemical shifts ($\delta$) are reported in parts per million (ppm) using the residual solvent ($CHCl_3$: $\delta$ = 7.26 ppm for $^{1}H$ NMR spectra and $CDCl_3$: $\delta$ = 77.00 ppm for $^{13}C$ NMR spectra) or TMS ($(CH_3)_4Si$: $\delta$ = 0 ppm for $^{1}H$ NMR spectra and $Me_4Si$: $\delta$ = 0 ppm for $^{13}C$ NMR spectra) as an internal standard. Multiplicities are given as s (singlet), d (doublet), t (triplet), q (quadruplet), m (multiplett). Coupling constants (*J*) are reported in Hz. HRMS analyses and accurate mass determinations were performed on a Thermo Scientific LTQ Orbitrap XL mass spectrometer using ESI ionisation and positive or negative mode by the analytical services (mass spectrometry lab of Prof. Dr. Stefan Schürch) from the Department of Chemistry, Biochemistry and Pharmaceutical Sciences (DCBP) of the University of Bern, Switzerland. HPLCs were measured on a Thermo-Scientific UltimMate 3000 HPLC with $H_2O$ + 0.1 % TFA and MeCN + 0.1 % TFA as eluents on a Acclaim ™ 120 C18 5 µm 120 Å (4.6 x 150 mm) column.

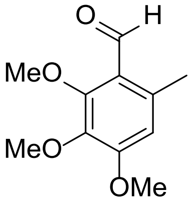

2,3,4-trimethoxy-6-methylbenzaldehyde
Chemical Formula: $C_{11}H_{14}O_4$

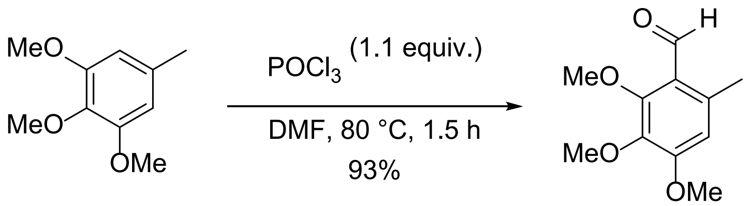

To a stirred solution of 3,4,5-trimethoxytoluene (10.047 g, 55.137 mol) in DMF (10 mL) $POCl_3$ (5.8 mL, 9.512 g, 62.036 mmol) was added dropwise over a period of 30 minutes through an addition funnel at 45 °C. Then the reaction mixture was warmed to 80 °C and stirred for 1.5 h. The mixture was cooled to 21 °C using an ice bath. Then the mixture was poured onto approximately 100 mL of wet ice. The pH was adjusted to ≈7 using aq. NaOH (ca. 100 mL 2 M NaOH). A mixture of ethyl acetate and dichloromethane was added to this solution. The layers were separated, and the aqueous phase was extracted with a mixture of ethyl acetate and dichloromethane. The combined organic phases were washed with brine (3x), dried over $MgSO_4$, filtered, and concentrated under reduced pressure. The title compound was obtained in 93 % yield (10.832 g, 51.525 mmol) as yellowish solid.

**$^1$H NMR** (300 MHz, $CDCl_3$) $\delta$ 10.40 (d, $J$ = 0.6 Hz, 1H), 6.50 (s, 1H), 3.97 (s, 3H), 3.91 (s, 3H), 3.85 (s, 3H), 2.56 (d, $J$ = 0.7 Hz, 3H). **$^{13}$C NMR** (75 MHz, $CDCl_3$) $\delta$ 191.17, 158.45, 157.77, 139.78, 138.24, 121.44, 110.51, 62.48, 61.09, 56.13, 21.89. **HRMS** (ESI) calculated for $[M+H]^+$ 211.0965, found 211.0959.

GA_216016.10.fid
PG05-001-02-03
Proton_ns32 CDCl3 /opt lochner 26

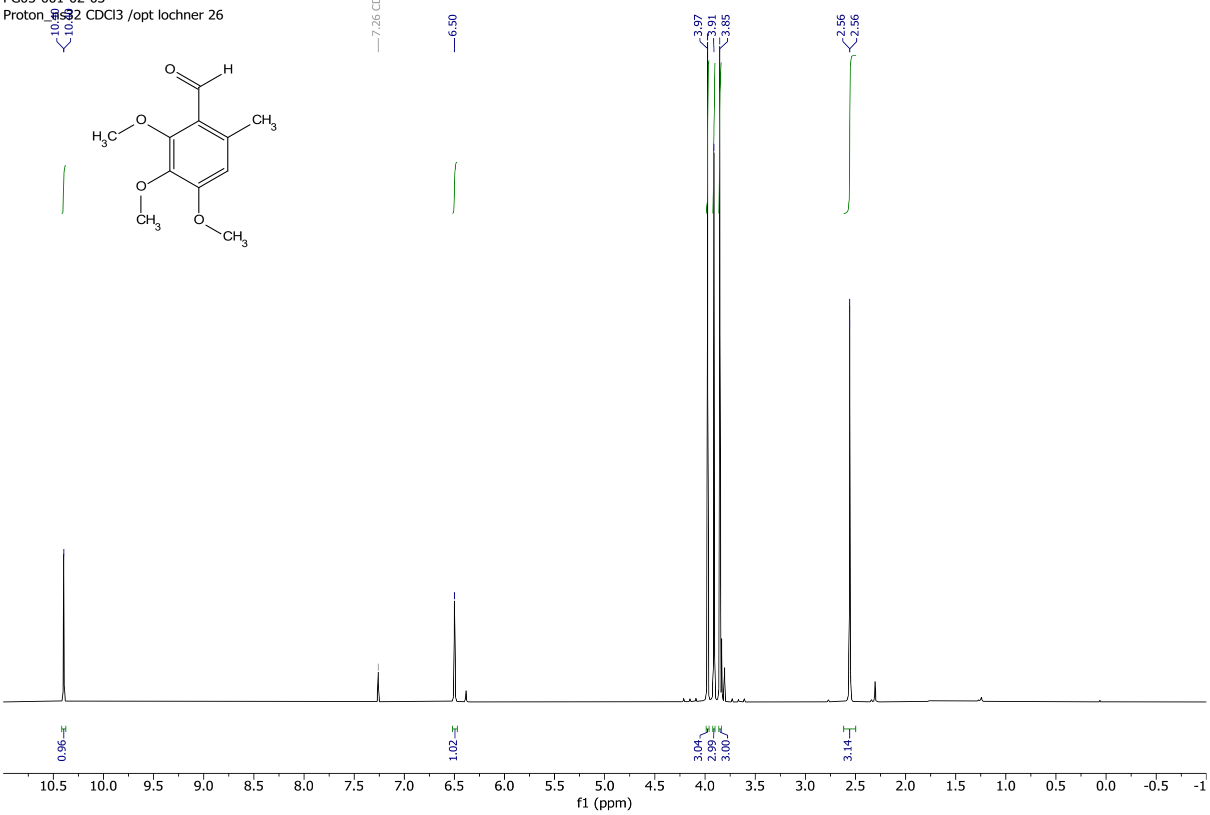

GA_216016.11.fid
PG05-001-02-03
Carbon_ns512 CDCl3 /opt lochner 26

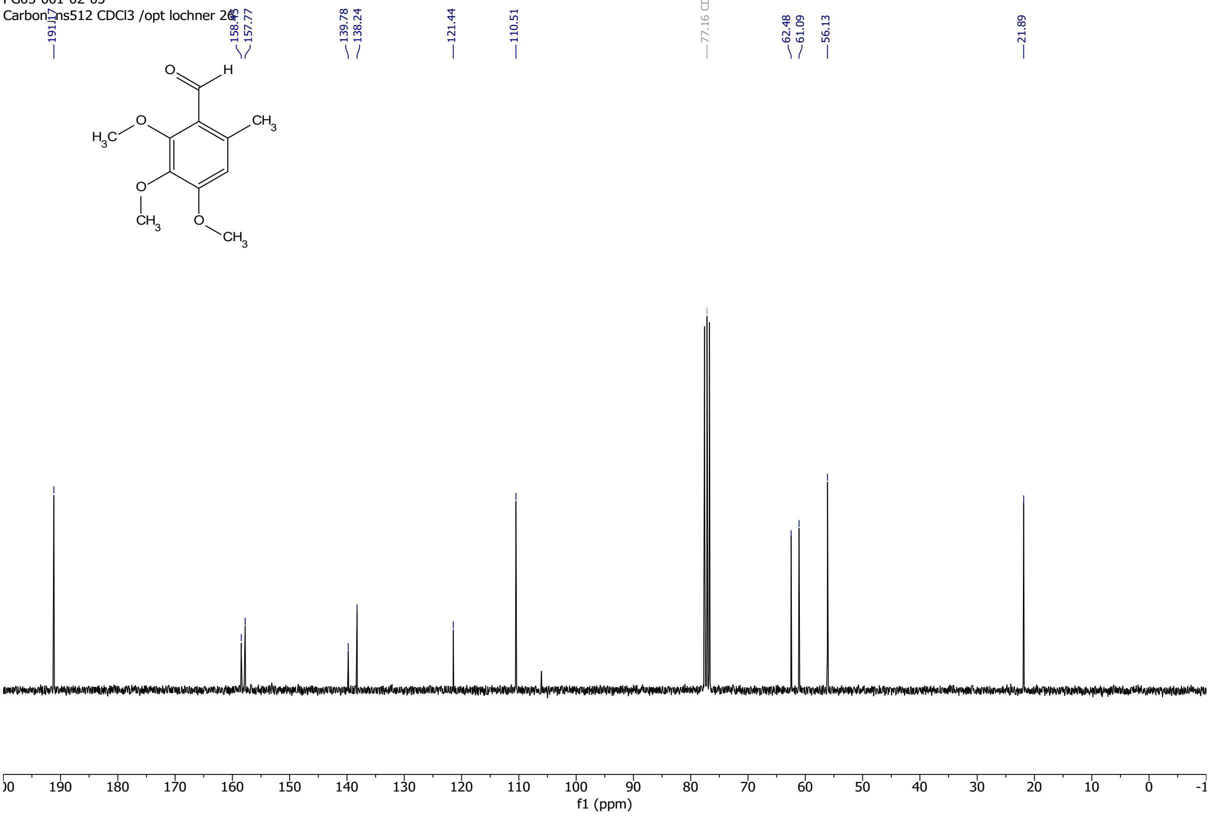

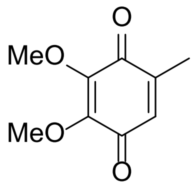

2,3-dimethoxy-5-methylcyclohexa-2,5-diene-1,4-dione
Chemical Formula: $C_9H_{10}O_4$

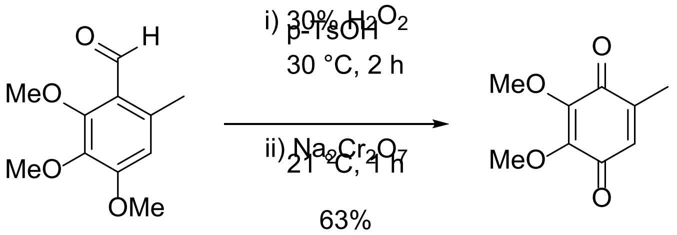

To a stirred solution of 2,3,4-trimethoxy-6-methylbenzaldehyde (1.0000 g, 4.7567 mmol) and *p*-TsOH monohydrate (0.4520 g, 2.3762 mmol) in methanol (5 mL) at 30 °C hydrogen peroxide (30% in water, 0.6 mL, 6.4603 mmol) was added slowly. After stirring for 2 h at 30 °C to the orange mixture sodium dichromate (0.3931 g, 1.3188 mmol) was added. This immediately led to the formation of a greenish-black solution. The mixture was stirred for 1 h at room temperature and its colour changed to brown. Petrol ether (bp 50-70 °C) was added, the layers separated, which resulted in an orange-coloured organic phase. The aqueous phase was extracted with petrol ether (bp 50-70 °C). The combined organic layers were washed with brine, dried over $MgSO_4$, filtered, and concentrated under reduced pressure. The title compound was obtained in 63% yield (0.54240 g, 2.9773 mmol) as a red solid. The physical appearance and the spectral data were in accordance with literature values.

**$^1$H NMR** (300 MHz, $CDCl_3$) $\delta$ 6.42 (d, *J* = 1.6 Hz, 1H), 4.01 (s, 3H), 3.98 (s, 3H), 2.03 (d, *J* = 1.7 Hz, 3H).
**$^{13}$C NMR** (75 MHz, $CDCl_3$) $\delta$ 184.54, 184.31, 145.15, 144.96, 144.17, 131.41, 61.38, 61.31, 15.59.
**HRMS** (ESI) calculated for $[M+H]^+$ 183.0652, found 183.0650.

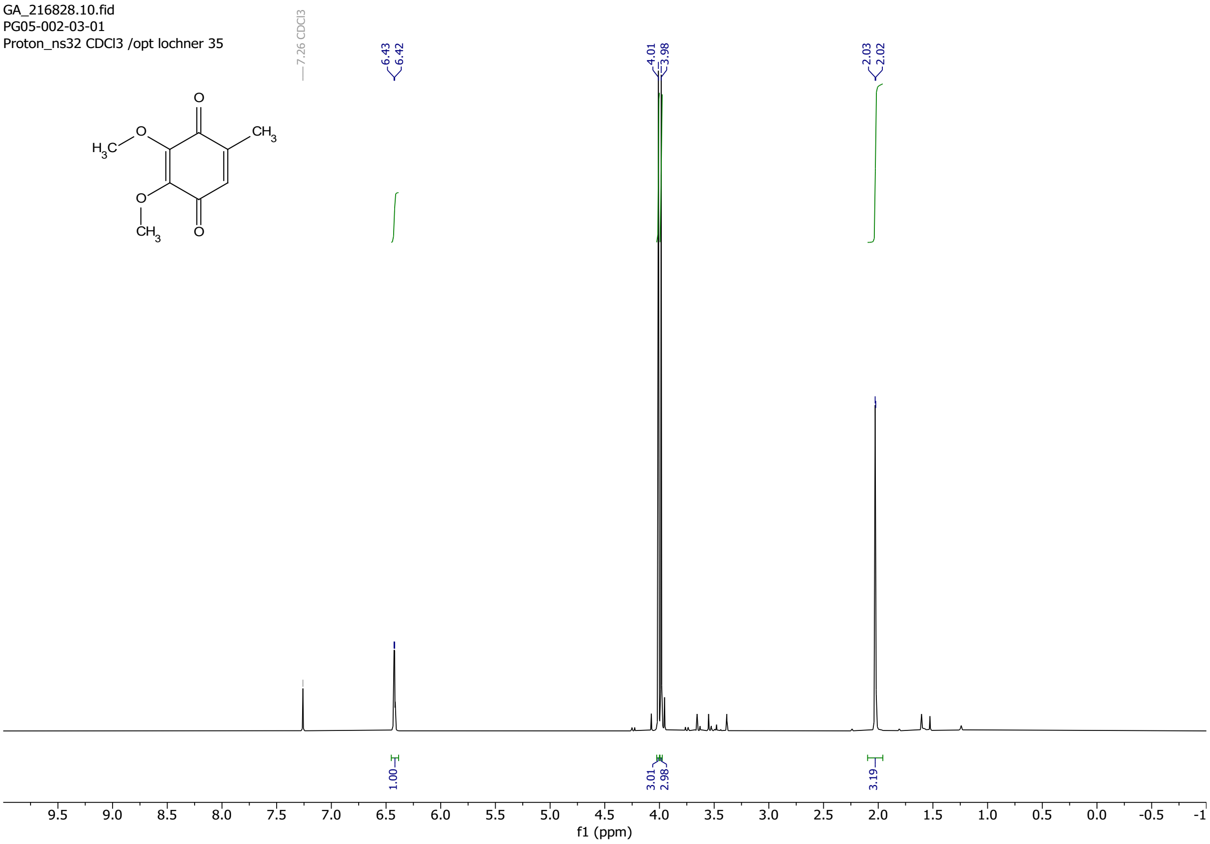
GA_216828.10.fid
PG05-002-03-01
Proton_ns32 CDCl3 /opt lochner 35
7.26 CDCl3
6.43
6.42
4.01
3.98
2.03
2.02
1.00
3.01
2.98
3.19
f1 (ppm)

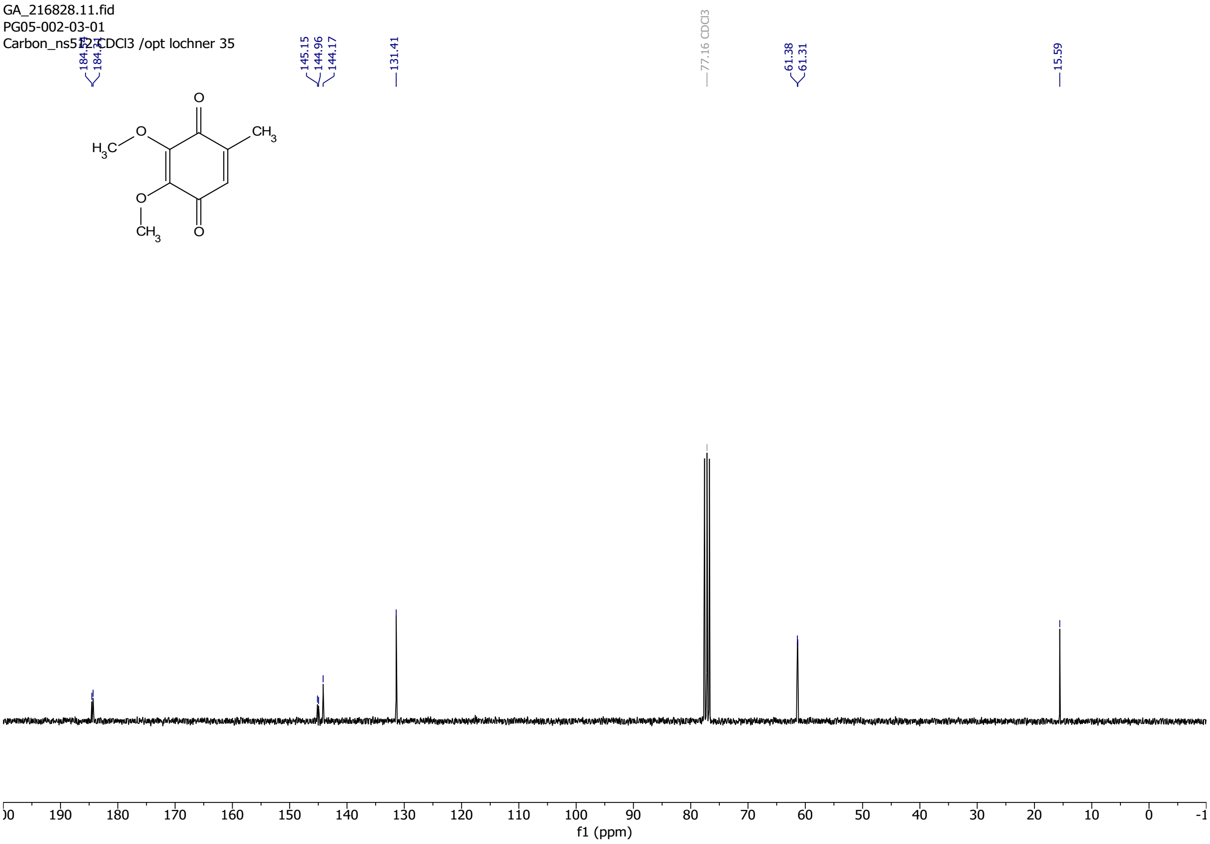
GA_216828.11.fid
PG05-002-03-01
145.15
144.96
144.17
131.41
77.16 CDCl3
61.38
61.31
15.59
f1 (ppm)

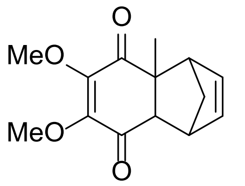

6,7-dimethoxy-4a-methyl-1,4,4a,8a-tetrahydro-1,4-methanonaphthalene-5,8-dione
Chemical Formula: $C_{14}H_{16}O_4$

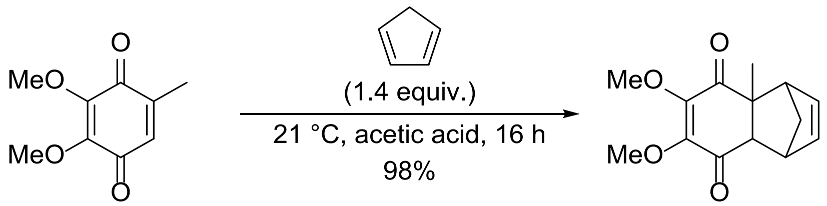

To a stirred solution of dimethoxy benzoquinone (3.34458 g, 18.3592 mmol) in acetic acid (8 mL) freshly distilled cyclopentadiene (2.2 mL, 1.76 g, 26.6251 mmol) was added and the red mixture was stirred under an argon atmosphere at 21 °C for 16 h. The red colour slowly changed into a faint orange. After 16 h of reaction time, the mixture was concentrated under reduced pressure, then taken up in dichloromethane and concentrated onto silica gel. The crude was purified by flash column chromatography (cyclohexane/ ethyl acetate gradient from 1:0 to 7:3). The title compound was obtained in 98% yield (4.45771 g, 17.9540 mmol) as a yellow oil. The physical appearance and the spectral data were in accordance with literature values.

**$^1$H NMR** (300 MHz, $CDCl_3$) $\delta$ 6.08 (ddd, $J$ = 43.7, 5.7, 2.9 Hz, 2H), 3.94 (s, 3H), 3.92 (s, 3H), 3.42 (p, $J$ = 1.6 Hz, 1H), 3.14 – 3.03 (m, 1H), 2.82 (d, $J$ = 3.9 Hz, 1H), 1.69 – 1.51 (m, 2H), 1.48 (s, 3H). **$^{13}$C NMR** (75 MHz, $CDCl_3$) $\delta$ 198.62, 195.02, 150.73, 150.67, 138.30, 134.64, 60.79 (2 C), 57.21, 53.54, 52.69, 48.97, 46.48, 26.65. **HRMS** (ESI) calculated for $[M+Na]^+$ 271.0941, found 271.0935.

GA_216827.10.fid
PG05-004-01-01
Proton_ns32 CDCl3 /opt lochner 34

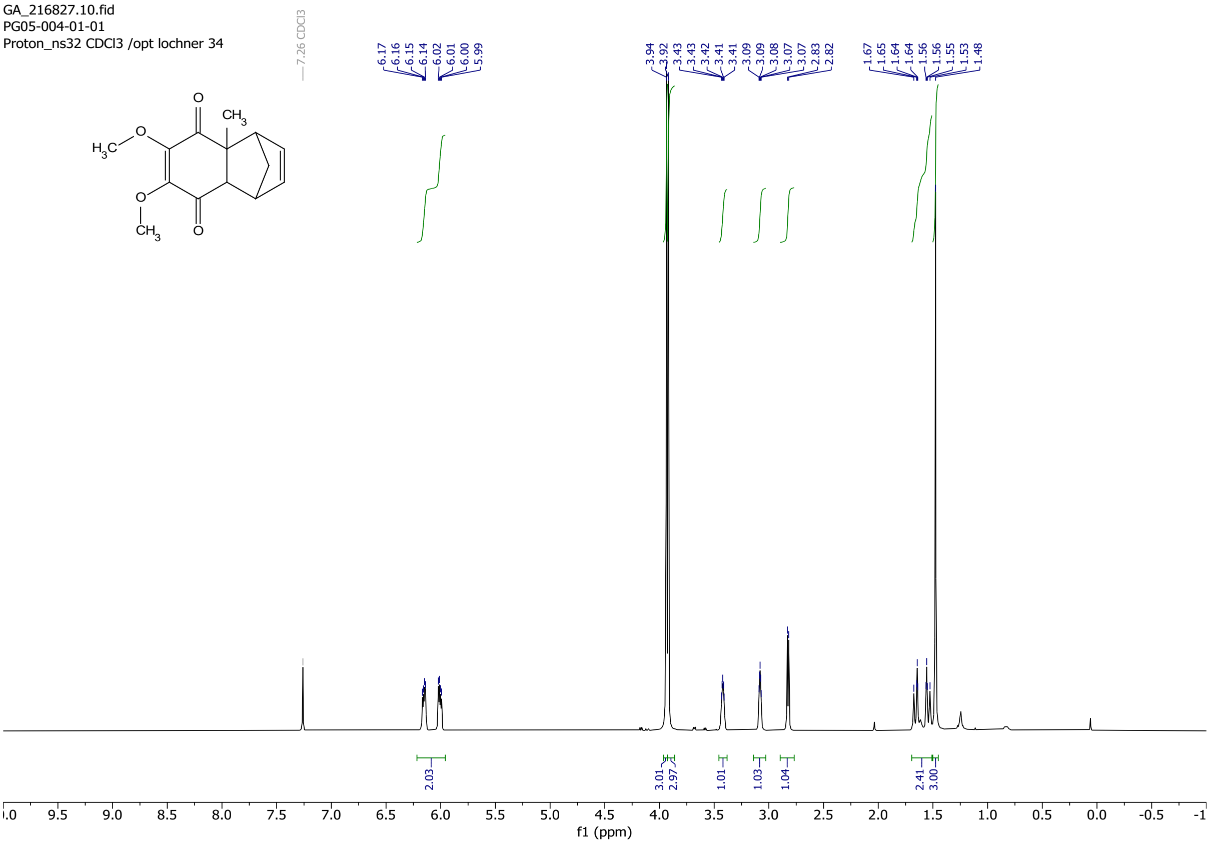

GA_216827.11.fid
PG05-004-01-01
Carbon_ns512 CDCl3 /opt lochner 34

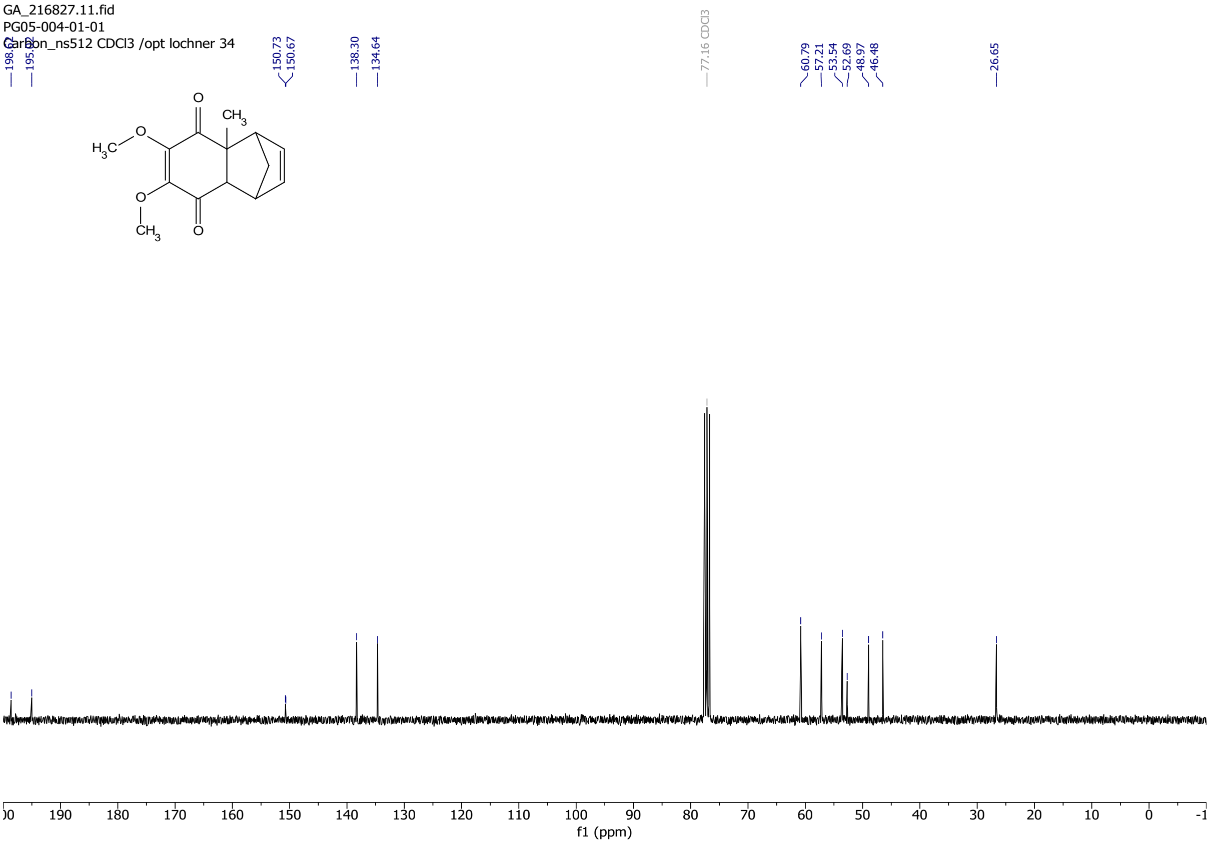

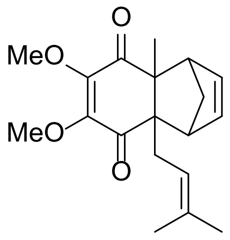

6,7-dimethoxy-4a-methyl-8a-(3-methylbut-2-en-1-yl)-1,4,4a,8a-tetrahydro-1,4-methanonaphthalene-5,8-dione
Chemical Formula: $C_{19}H_{24}O_4$

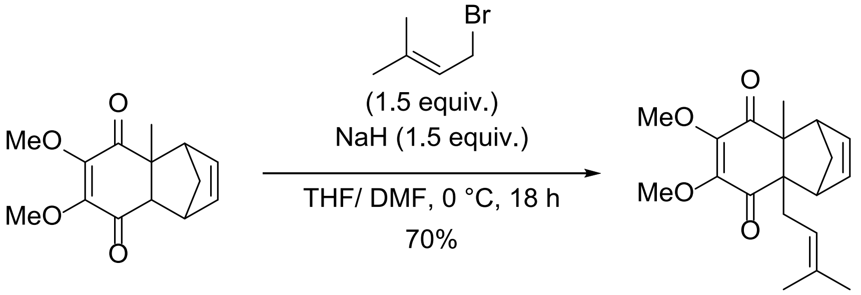

To a stirred solution of 6,7-dimethoxy-4a-methyl-1,4,4a,8a-tetrahydro-1,4-methanonaphthalene-5,8-dione (0.2638 g,1.0625 mmol) in dry THF (2 mL) at 0 °C NaH (60 % dispersion in parrafinum, 0.0498 g, 2.0759 mmol) was added. The mixture was stirred for 60 minutes at 0 °C giving a brown slurry. After 1 h stirring at 0 °C, 1-bromo-3-methyl-2-butene (1.0 mL, 1.29 g, 8.6559 mmol) was added dropwise. The mixture gradually turned into a yellowish slurry. It was left stirring for 16 h at room temperature. Then DMF (2 mL) was added, and the mixture was stirred for another 1 h. The reaction was carefully quenched with water. The aqueous phase was extracted with ethyl acetate. The combined organic layers were dried over $MgSO_4$, filtered, and concentrated under reduced pressure. The crude was taken up in dichloromethane and concentrated onto silica gel. The crude was purified by flash column chromatography (cyclohexane/ ethyl acetate gradient from 1:0 to 3:2). The title compound was obtained in 70 % yield (0.23552 g, 0.7444 mmol) as yellow oil.

**$^1$H NMR** (300 MHz, $CDCl_3$) δ 6.05 (dt, *J* = 2.6, 1.2 Hz, 2H), 5.08 (tdd, *J* = 6.3, 2.9, 1.4 Hz, 1H), 3.91 (s, 3H), 3.87 (s, 3H), 3.05 (dq, *J* = 23.6, 1.8 Hz, 2H), 2.80 – 2.33 (m, 2H), 1.81 – 1.73 (m, 1H), 1.67 (s, 3H), 1.58 (s, 3H), 1.50 (s, 3H), 1.48 – 1.40 (m, 2H). **$^{13}$C NMR** (75 MHz, $CDCl_3$) *δ* 199.02, 198.41, 150.96, 149.28, 138.11, 137.34, 134.61, 119.98, 60.46, 60.11, 59.44, 56.22, 54.57, 53.27, 43.60, 36.25, 26.18, 23.53, 18.06. **HRMS** (ESI) calculated for $[M+H]^+$ 317.1747, found 317.1750.

GA_216936.10.fid
PG05-004-02-01
Proton_ns32 CDCl3 /opt kuepfer 14

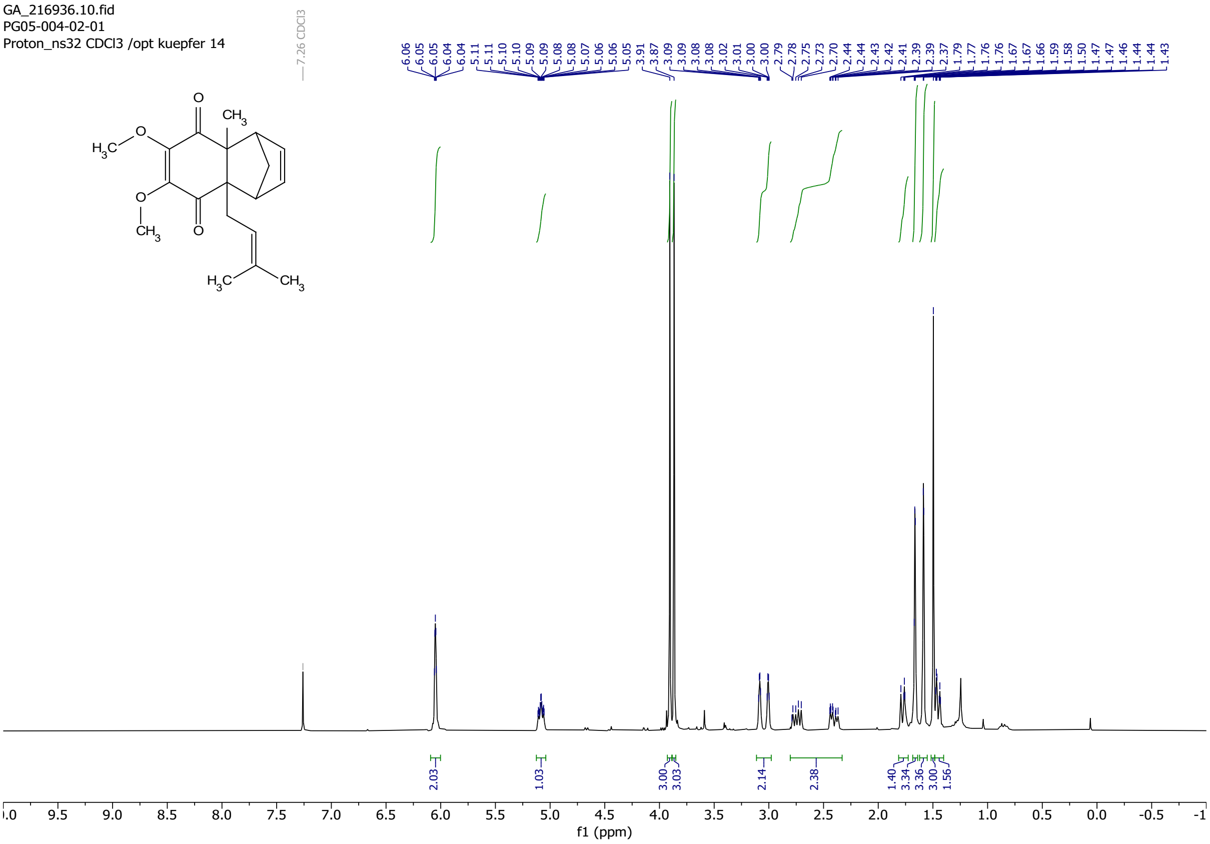

GA_216936.11.fid
Carbon_ns512 CDCl3 /opt lochner 14

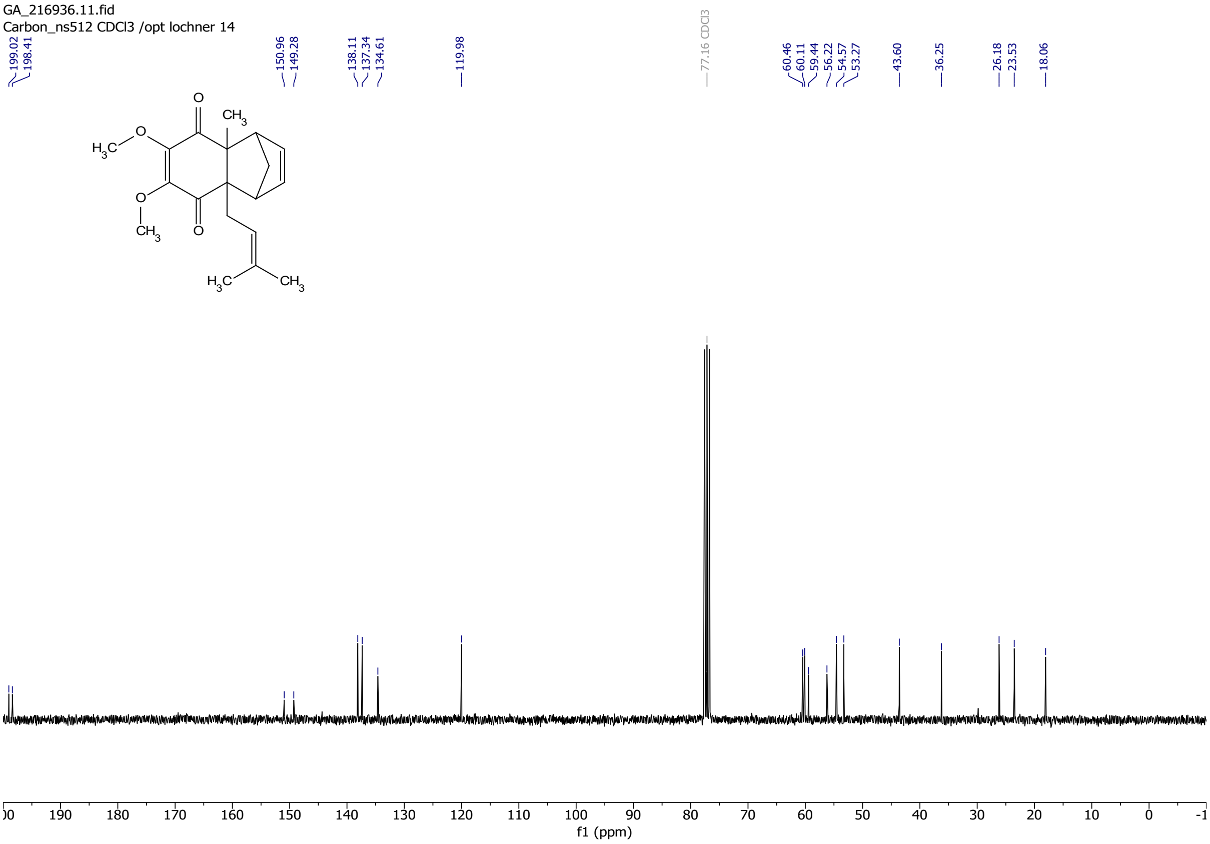

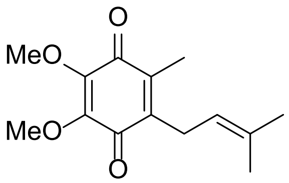

2,3-dimethoxy-5-methyl-6-(3-methylbut-2-en-1-yl)cyclohexa-2,5-diene-1,4-dione
Chemical Formula: $C_{14}H_{18}O_4$

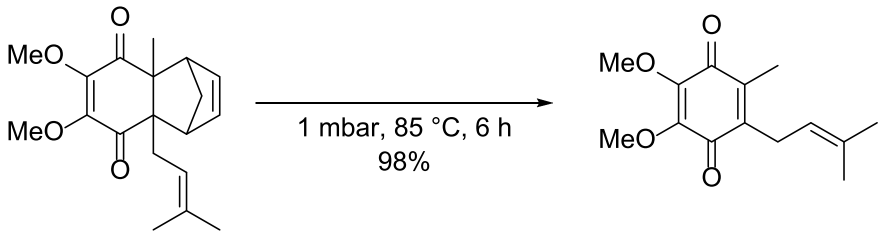

6,7-dimethoxy-4a-methyl-8a-(3-methylbut-2-en-1-yl)-1,4,4a,8a-tetrahydro-1,4-methanonaphthalene-5,8-dione (0.22311 g, 0.7052 mmol) was heated to 85 °C under reduced atmosphere (~1 mbar). The mixture was protected from light with tinfoil. The orange oil gradually changed its colour to an intense red. After 6 h the compound was cooled to room temperature. The crude was purified by reverse phase flash column chromatography (C18, $H_20$ + 0.1 % TFA/ MeCN + 0.1 % TFA gradient from 1:0 to 0:1). The title compound was obtained in 98 % yield (0.17247 g, 0.6890 mmol) as a red oil.

**$^1$H NMR** (300 MHz, $CDCl_3$) $\delta$ 4.93 (tp, *J* = 7.1, 1.4 Hz, 1H), 3.99 (s, 3H), 3.98 (s, 3H), 3.17 (d, *J* = 7.1 Hz, 2H), 2.02 (s, 3H), 1.74 (s, 3H), 1.67 (d, *J* = 1.4 Hz, 3H). **$^{13}$C NMR** (75 MHz, $CDCl_3$) $\delta$ 184.95, 184.11, 144.55, 144.39, 141.76, 138.95, 134.16, 119.15, 61.30, 25.87, 25.58, 18.11, 12.08. **HRMS** (ESI) calculated for $[M+H]^+$ 251.1278, found 251.1287.

**Purity (HPLC)**: 93 % (at 214 nm), 96 % (at 254 nm).

GA_217296.10.fid
PG05-005-02-01
Proton_ns32 CDCl3 /opt lochner 50

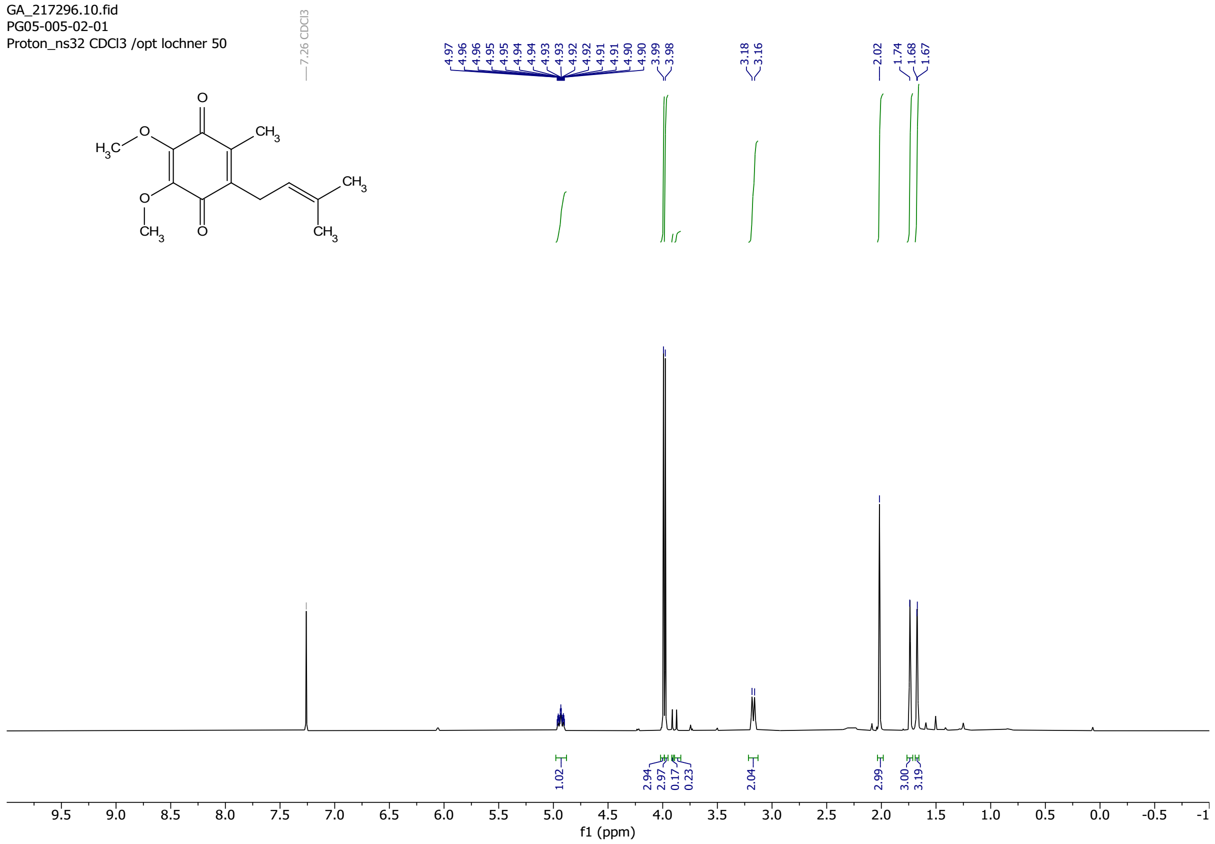

GA_217158.11.fid
PG05-005-01-02
Carbon_ns512 CDCl3 /opt lochner 52

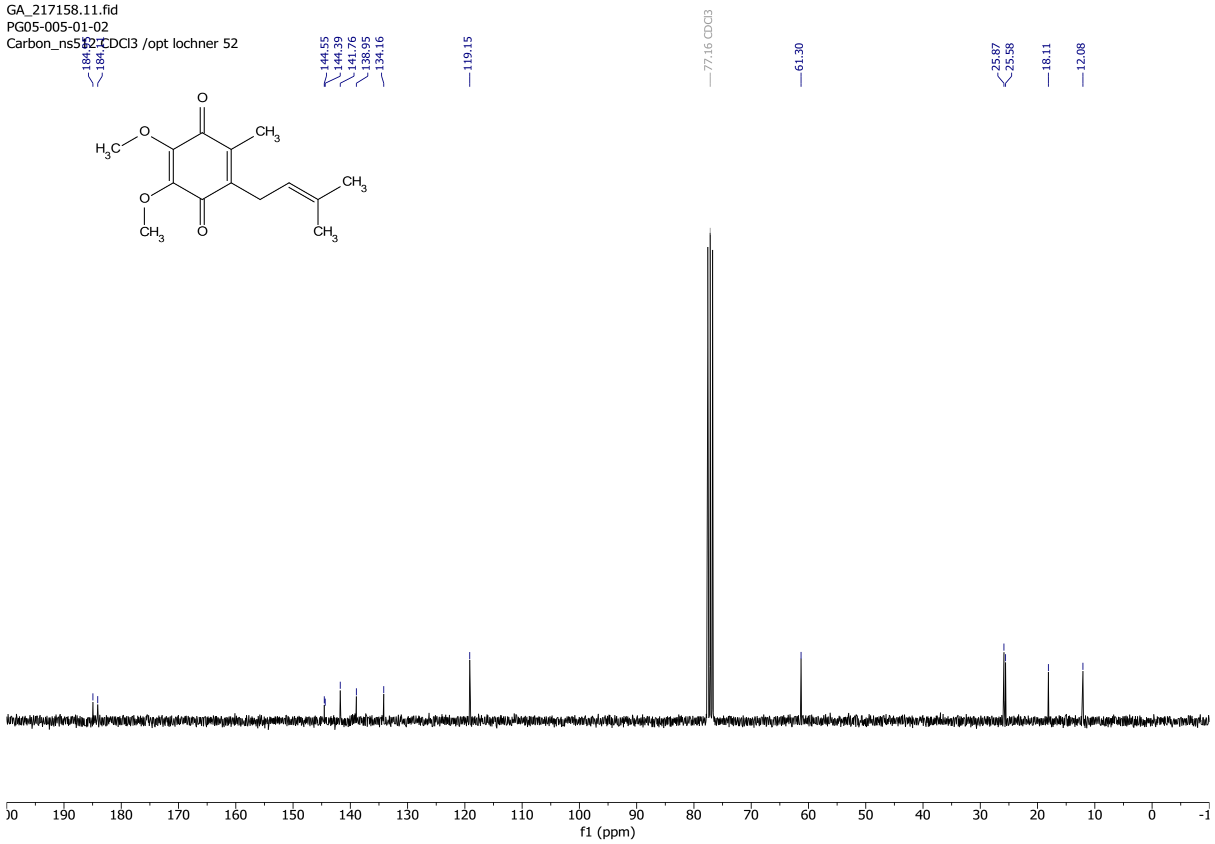

#### Chromatogram

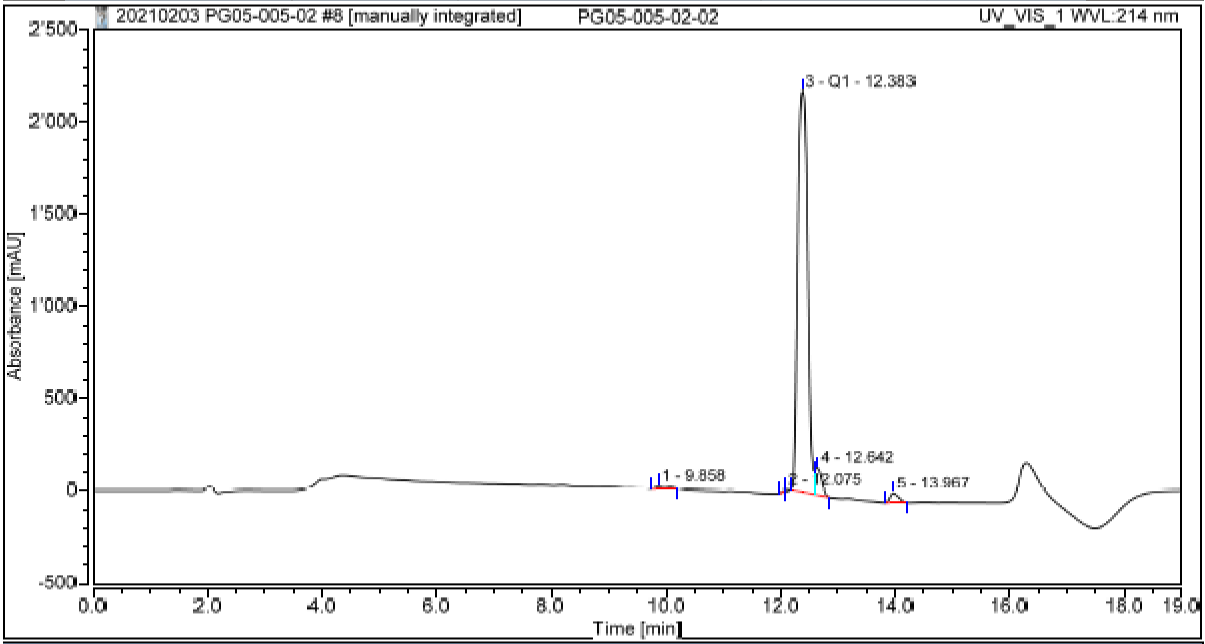

#### Integration Results

| No. | Peak Name | Retention Time min | Area mAU*min | Height mAU | Relative Area % | Relative Height % | Amount |
|---|---|---|---|---|---|---|---|
| n.a. | None | n.a. | n.a. | n.a. | n.a. | n.a. | n.a. |
| 1 | | 9.858 | 2.669 | 11.320 | 0.57 | 0.47 | n.a. |
| 2 | | 12.075 | 0.388 | 10.664 | 0.08 | 0.45 | n.a. |
| 3 | Q1 | 12.383 | 435.115 | 2173.803 | 93.20 | 90.96 | n.a. |
| 4 | | 12.642 | 20.603 | 146.004 | 4.41 | 6.11 | n.a. |
| 5 | | 13.967 | 8.110 | 47.994 | 1.74 | 2.01 | n.a. |
| **Total:** | | | **466.885** | **2389.784** | **100.00** | **100.00** | |

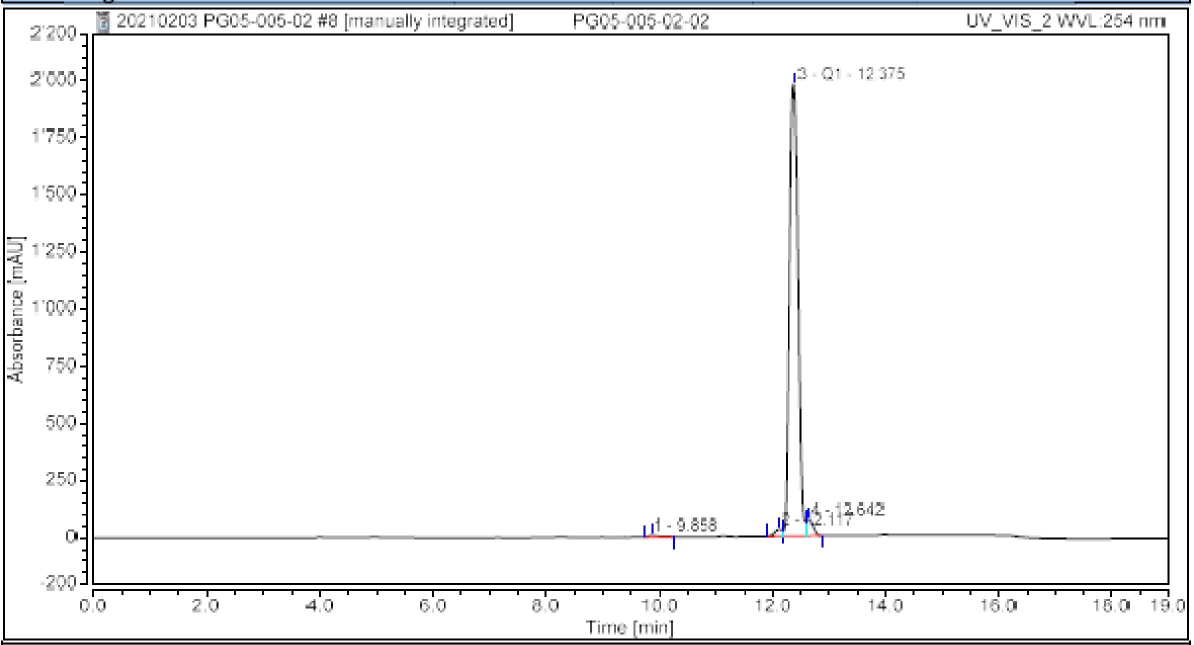

Integration Results

| No. | Peak Name | Retention Time min | Area mAU*min | Height mAU | Relative Area % | Relative Height % | Amount |
|---|---|---|---|---|---|---|---|
| n.a. | None | n.a. | n.a. | n.a. | n.a. | n.a. | n.a. |
| 1 | | 9.858 | 1.657 | 7.522 | 0.45 | 0.36 | n.a. |
| 2 | | 12.117 | 4.343 | 29.584 | 1.18 | 1.42 | n.a. |
| 3 | Q1 | 12.375 | 352.042 | 1972.030 | 95.64 | 94.81 | n.a. |
| 4 | | 12.642 | 10.060 | 70.754 | 2.73 | 3.40 | n.a. |
| **Total:** | | | **368.102** | **2079.890** | **100.00** | **100.00** | |

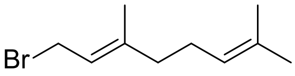

(*E*)-1-bromo-3,7-dimethylocta-2,6-diene
Chemical Formula: $C_{10}H_{17}Br$

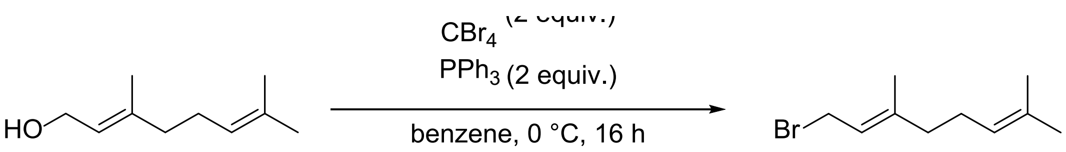

To a stirred solution of geraniol (2.501 g, 16.2136 mmol) and tetrabromomethane (10.818 g, 32.6027 mmol) in benzene (50 mL) at 0 °C triphenylphosphine (8.533 g, 32.5327 mmol) was added in portions. The mixture was stirred at 0 °C. It was stirred for another 16 h while slowly warming to 20 °C. To the mixture was added petrolether (bp 50-70 °C). This led to a brown solid. This solid was filtered off, and the filtrate was concentrated under reduced pressure. To the resulting clear colourless liquid residue petrolether (bp 50-70 °C) was added. This led to a lot of white precipitate. This mixture was filtered through a pad of celite. The filtrate was concentrated under reduced pressure. The crude was distilled twice in a Kugelrohr apparatus at ~1mbar and 120 °C. The title compound was obtained in 83% yield (2.93063 g, 13.4959 mmol) of a clear colorless low viscosity oil.

**$^{1}$H NMR** (300 MHz, $CDCl_3$) *δ* 5.60 – 5.47 (m, 1H), 5.09 – 5.05 (m, 1H), 4.03 (d, *J* = 8.4 Hz, 2H), 2.16 – 2.00 (m, 4H), 1.73 (d, *J* = 1.4 Hz, 3H), 1.68 (s, 3H), 1.60 (s, 3H). **$^{13}$C NMR** (75 MHz, $CDCl_3$) δ 143.78, 132.17, 123.68, 120.68, 39.68, 29.86, 26.35, 25.82, 17.86, 16.12. **HRMS** (ESI) calculated for $[M+H]^+$, found.

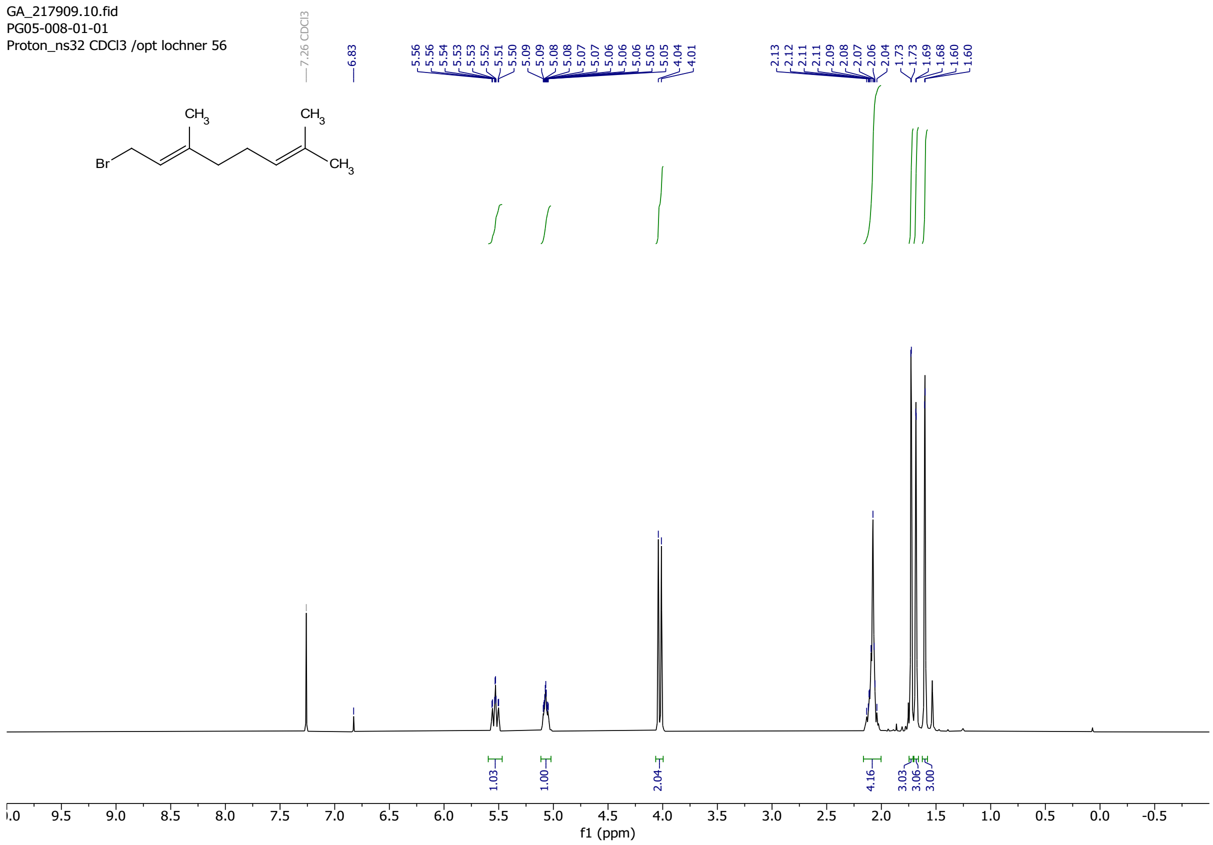

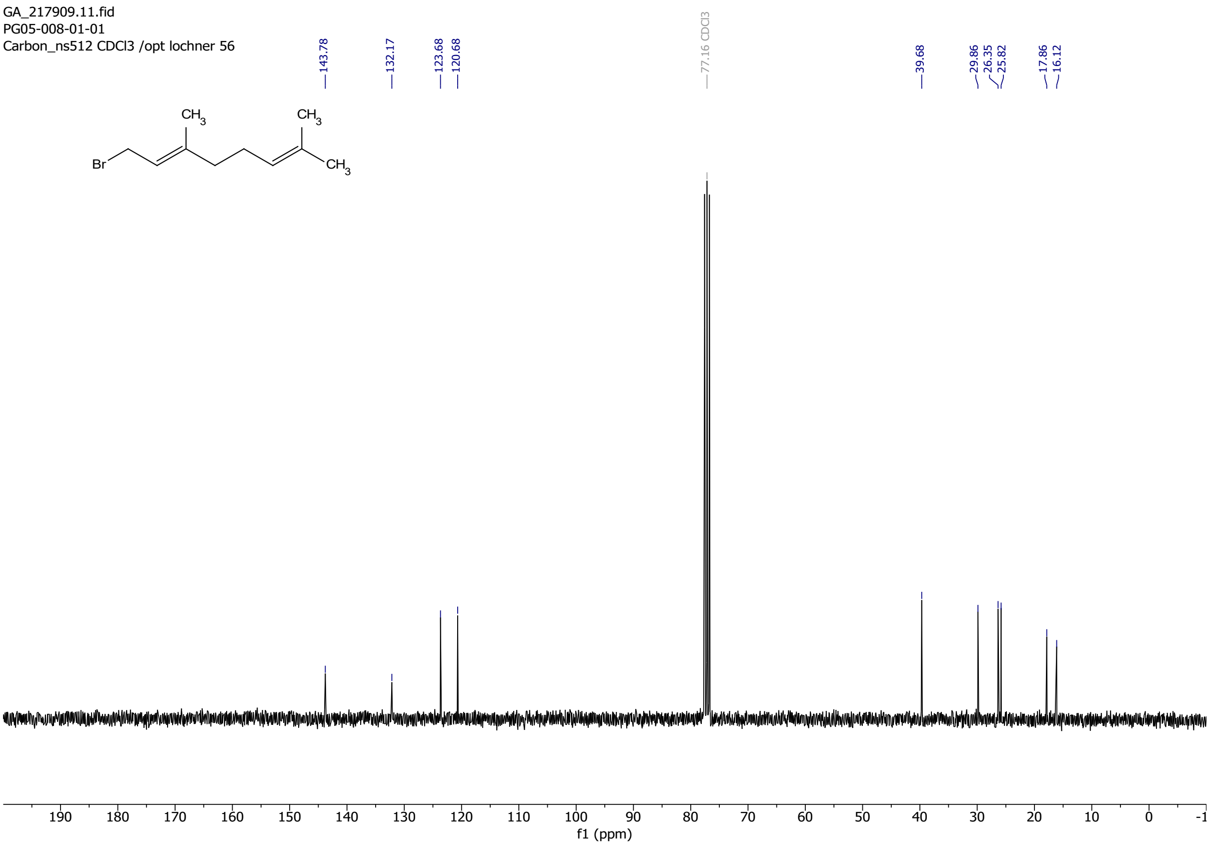

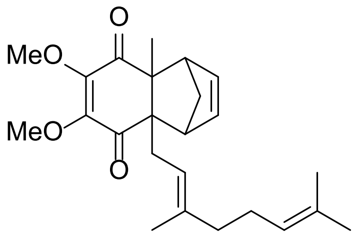

(*E*)-4a-(3,7-dimethylocta-2,6-dien-1-yl)-6,7-dimethoxy-8a-methyl-1,4,4a,8a-tetrahydro-1,4-methanonaphthalene-5,8-dione

Chemical Formula: $C_{24}H_{32}O_4$

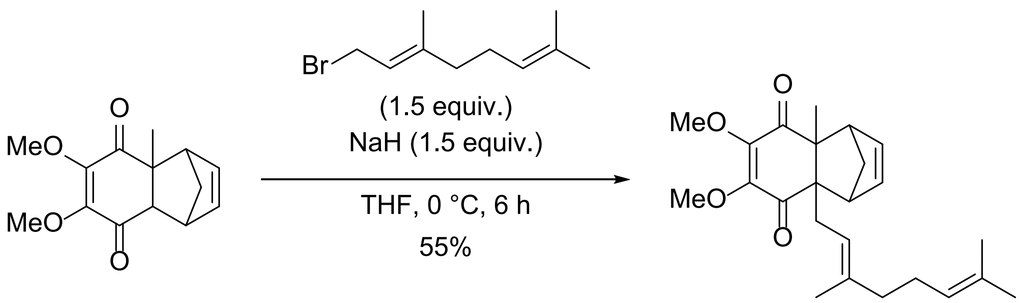

To a stirred solution of 6,7-dimethoxy-4a-methyl-1,4,4a,8a-tetrahydro-1,4-methanonaphthalene-5,8-dione (1.0099 g, 4.0676 mmol) in dry THF (5 mL) at 0 °C NaH (60 % dispersion in parrafinum, 0.290 g, 7.2530 mmol) was added. The mixture was stirred for 30

minutes at 0 °C, resulting in a white suspension. To this was added (*E*)-1-bromo-3,7-dimethylocta-2,6-diene (1.497 g, 6.8939 mmol) and DMF (1 mL). The mixture was slowly warmed to 21 °C and stirred for 16 h. The yellow solution was carefully quenched with water. The aqueous phase was extracted with ethyl acetate. The combined organic layers were washed with brine, dried over $MgSO_4$, filtered, and concentrated under reduced pressure to give a brown oil. The oil was taken up in dichloromethane and concentrated onto silica gel. The crude was purified by flash column chromatography (cyclohexane/ ethyl acetate gradient from 1:0 to 1:1). The title compound was obtained in 55 % yield (0.85634 g, 2.2271 mmol) as orange oil.

**$^1$H NMR** (300 MHz, $CDCl_3$) *δ* 6.08 – 6.03 (m, 2H), 5.12 – 5.01 (m, 2H), 3.90 (s, 3H), 3.88 (s, 3H), 3.05 (dd, *J* = 23.9, 1.7 Hz, 2H), 2.75 (dd, *J* = 15.2, 7.3 Hz, 1H), 2.43 (dd, *J* = 14.5, 5.8 Hz, 1H), 2.05 – 1.96 (m, 4H), 1.79 (d, *J* = 9.4 Hz, 1H), 1.65 (s, 3H), 1.58 (s, 3H), 1.60 – 1.54 (m, 3H), 1.49 (s, 3H), 1.46 (dt, *J* = 9.5, 1.7 Hz, 1H). **$^{13}$C NMR** (75 MHz, $CDCl_3$) *δ* 199.04, 198.48, 150.96, 149.33, 138.10, 138.05, 137.34, 131.83, 124.08, 119.86, 60.45, 60.21, 59.47, 56.20, 54.65, 53.37, 43.65, 40.05, 36.28, 26.59, 25.81, 23.47, 17.81, 16.46. **HRMS** (ESI) calculated for $[M+Na]^+$ 407.2193, found 407.2198.

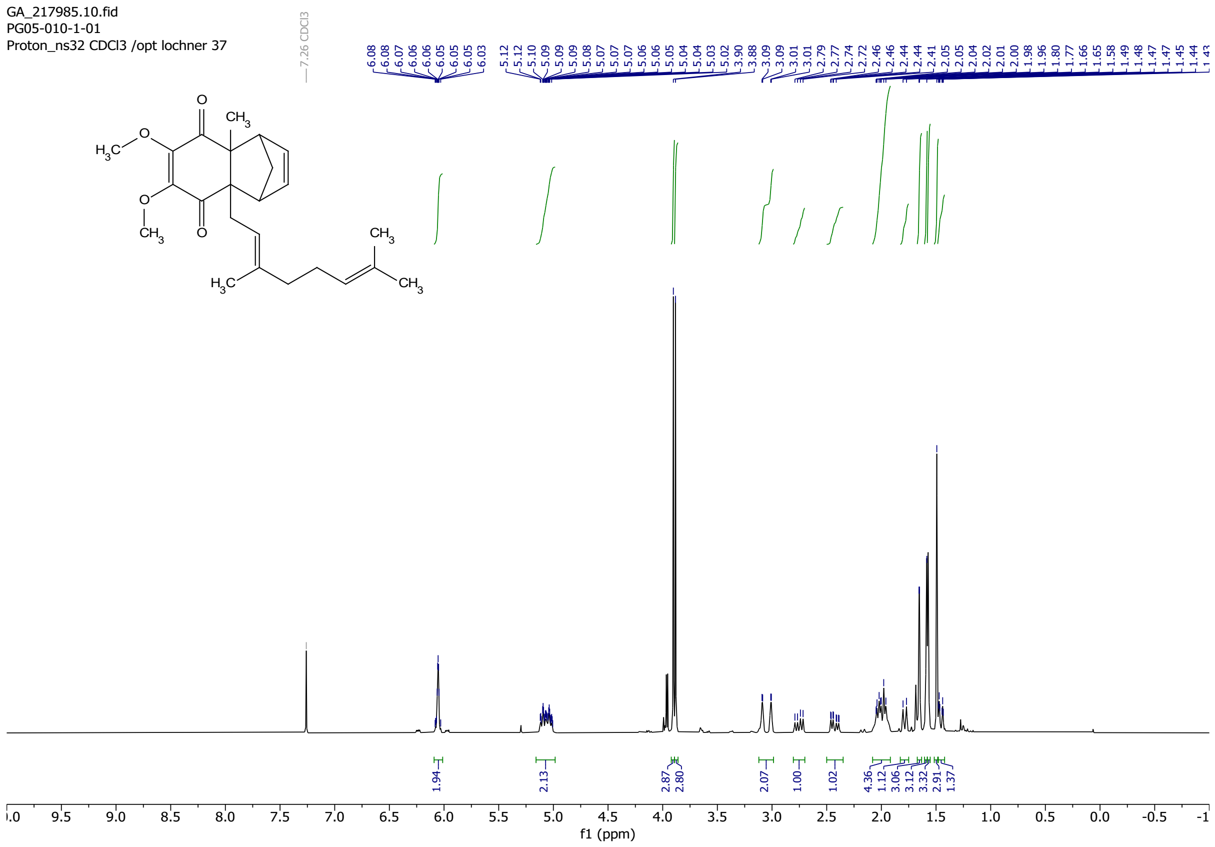

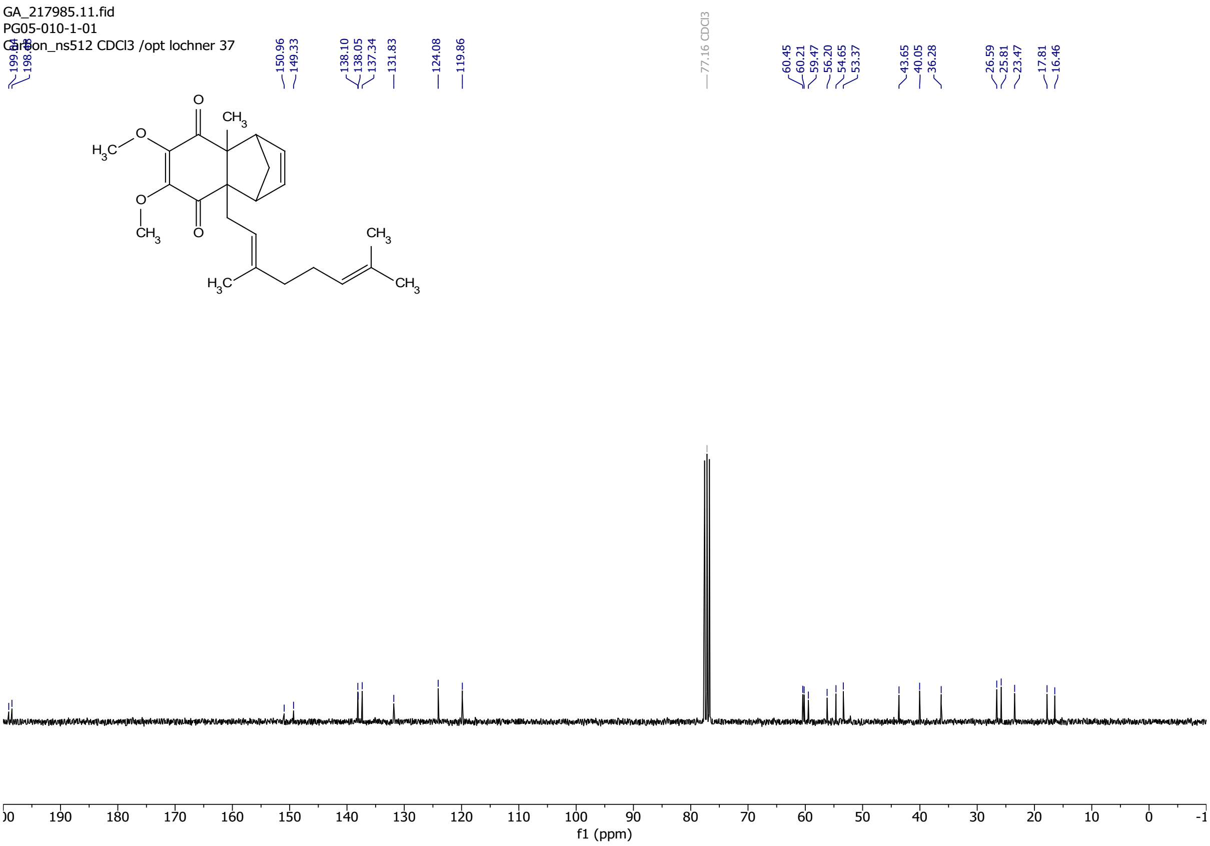

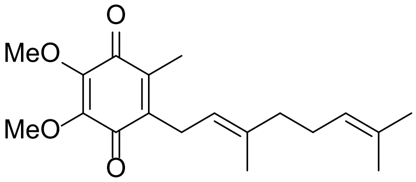

(*E*)-2-(3,7-dimethylocta-2,6-dien-1-yl)-5,6-dimethoxy-3-methylcyclohexa-2,5-diene-1,4-dione

Chemical Formula: $C_{19}H_{26}O_4$

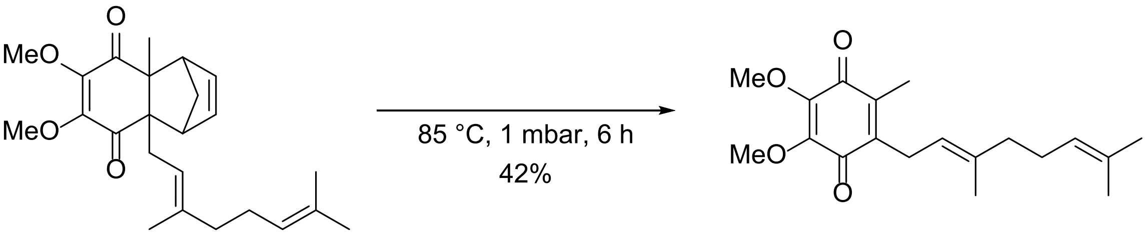

(*E*)-4a-(3,7-dimethylocta-2,6-dien-1-yl)-6,7-dimethoxy-8a-methyl-1,4,4a,8a-tetrahydro-1,4-methanonaphthalene-5,8-dione (0.82845 g, 2.1561 mmol) was heated to 85 °C at ~1 mbar for 6 h. After 6 h the red oil was taken up in dichloromethane and concentrated onto silica gel. The crude was purified by flash column chromatography (cyclohexane/ ethyl acetate 3:1). The isolated material was not pure, however. The mixture was taken up again in dichloromethane and concentrated onto silica. It was purified by reverse phase column chromatography (C18, $H_20$ + 0.1 % TFA/ MeCN + 0.1 % TFA gradient from 1:0 to 0:1). The title compound was obtained in 42 % yield (0.29119 g, 0.9145 mmol) as a red oil.

**$^1$H NMR** (300 MHz, $CDCl_3$) $\delta$ 5.08 – 4.97 (m, 1H), 4.93 (tq, $J$ = 7.1, 1.3 Hz, 1H), 3.99 (s, 3H), 3.98 (s, 3H), 3.21 – 3.16 (m, 1H), 2.09 – 1.92 (m, 7H), 1.73 (s, 3H), 1.65 (s, 3H), 1.57 (s, 3H). **$^{13}$C NMR** (75 MHz, $CDCl_3$) $\delta$ 184.90, 184.04, 141.82, 138.99, 137.63, 131.65, 129.14, 128.33, 124.11, 119.06, 61.25, 39.79, 26.62, 25.78, 25.40, 17.78, 16.39, 12.03. **HRMS** (ESI) calculated for $[M+Na]^+$ 319.1866, found 319.1873.

**Purity (HPLC)**: 91 % (at 214 nm), 95 % (at 254 nm).

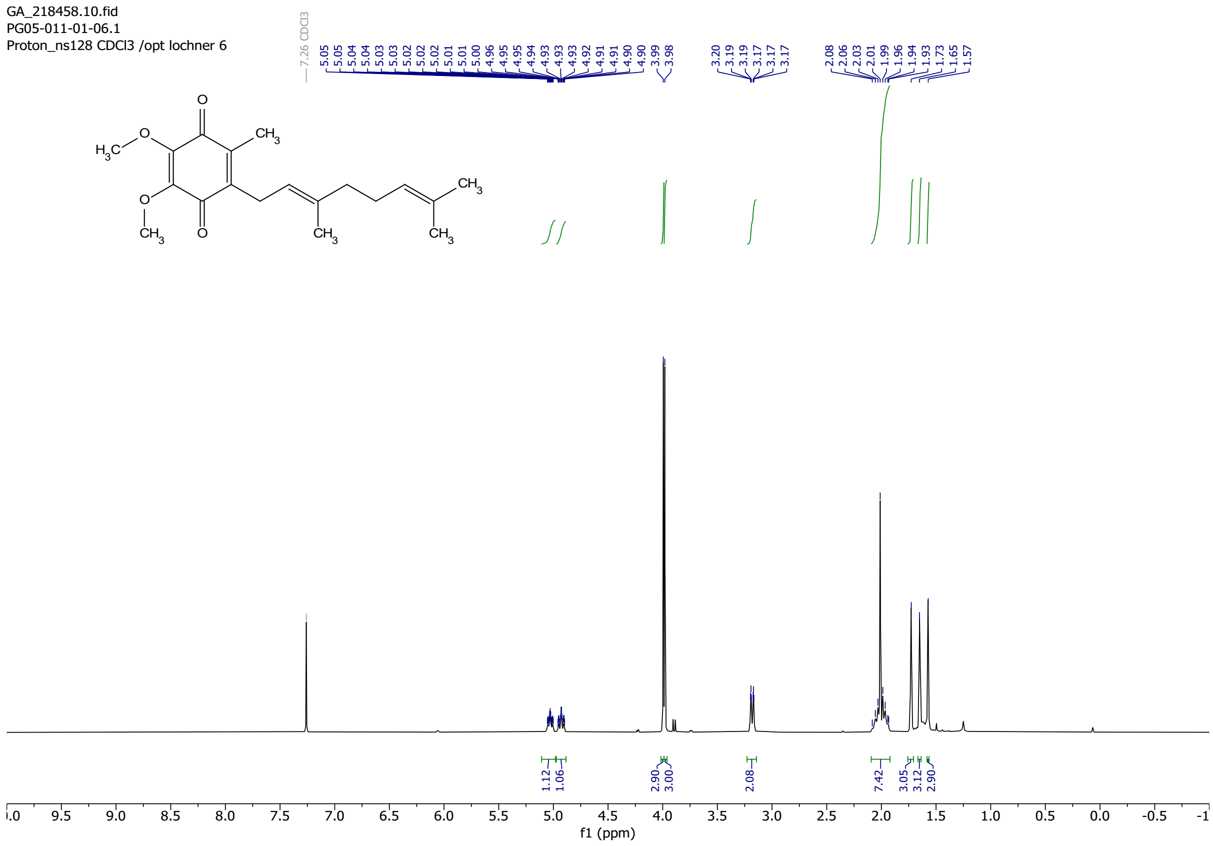

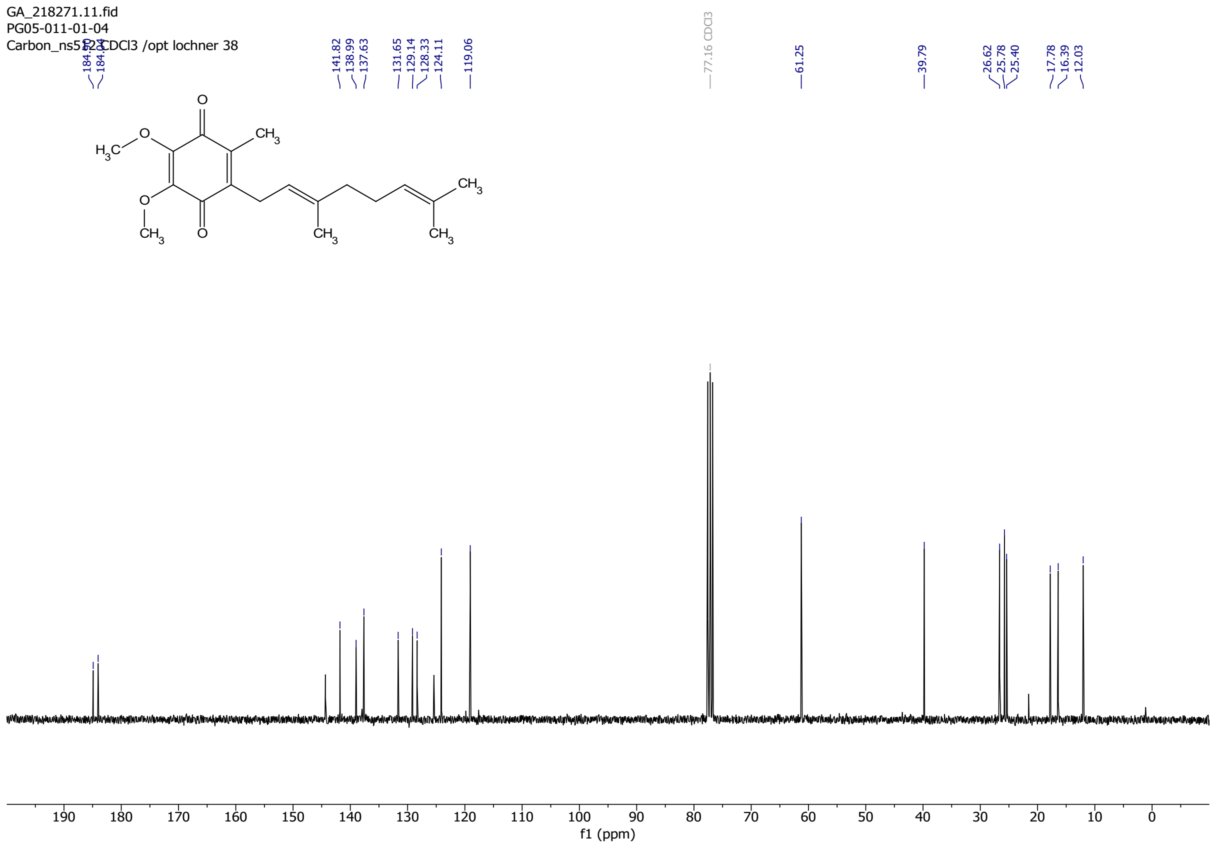
GA_218271.11.fid
PG05-011-01-04
Carbon_ns512 CDCl3 /opt lochner 38
184.90
184.04
141.82
138.99
137.63
131.65
129.14
128.33
124.11
119.06
77.16 CDCl3
61.25
39.79
26.62
25.78
25.40
17.78
16.39
12.03
O
CH3
H3C
O
O
CH3
O
CH3
CH3
CH3
190
180
170
160
150
140
130
120
110
100
90
80
70
60
50
40
30
20
10
0
f1 (ppm)

### Chromatogram and Results

#### Injection Details

| | | | |
|---|---|---|---|
| *Injection Name:* | **PG05-011-01-05** | *Run Time (min):* | **19.00** |
| *Vial Number:* | **BB2** | *Injection Volume:* | **2.50** |
| *Injection Type:* | **Calibration Standard** | *Channel:* | **UV_VIS_1** |
| *Calibration Level:* | | *Wavelength:* | **214** |
| *Instrument Method:* | **IM_General0%** | *Bandwidth:* | **4** |
| *Processing Method:* | **Q10** | *Dilution Factor:* | **1.0000** |
| *Injection Date/Time:* | **25.Feb.21 12:52** | *Sample Weight:* | **1.0000** |

#### Chromatogram

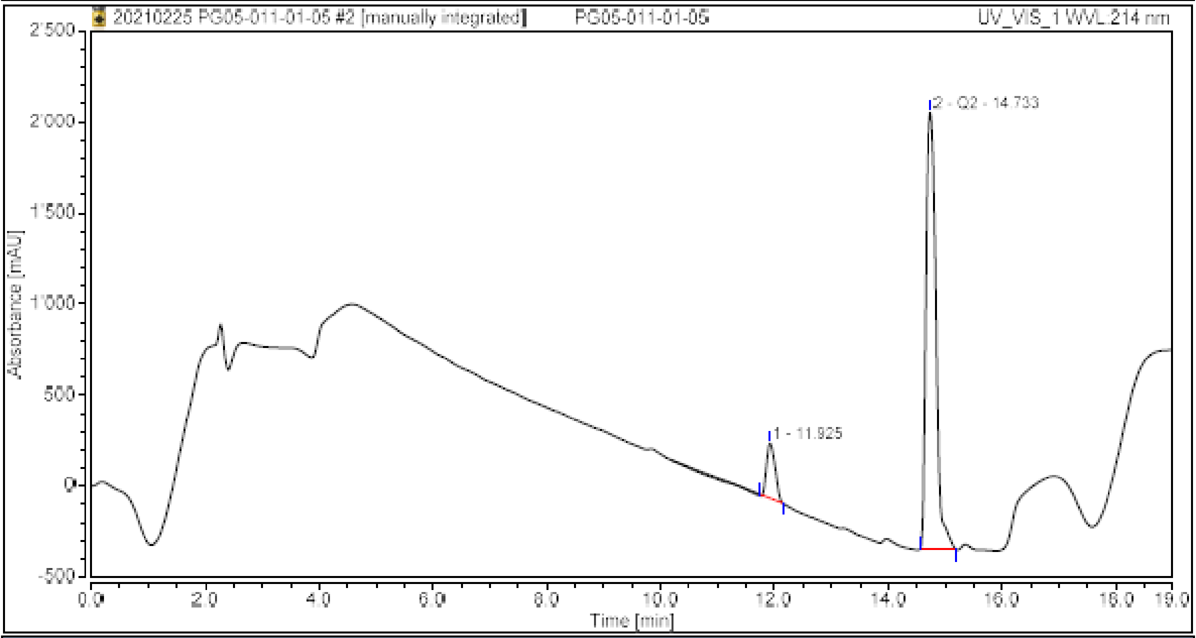

#### Integration Results

| No. | Peak Name | Retention Time<br>min | Area<br>mAU*min | Height<br>mAU | Relative Area<br>% | Relative Height<br>% | Amount |
|---|---|---|---|---|---|---|---|
| 1 | | 11.925 | 51.696 | 302.187 | 9.41 | 11.18 | n.a. |
| 2 | Q2 | 14.733 | 497.496 | 2399.663 | 90.59 | 88.82 | n.a. |
| **Total:** | | | **549.192** | **2701.850** | **100.00** | **100.00** | |

### Chromatogram and Results

#### Injection Details

| | | | |
|---|---|---|---|
| *Injection Name:* | **PG05-011-01-05** | *Run Time (min):* | **19.00** |
| *Vial Number:* | **BB2** | *Injection Volume:* | **2.50** |
| *Injection Type:* | **Calibration Standard** | *Channel:* | **UV_VIS_2** |
| *Calibration Level:* | | *Wavelength:* | **254** |
| *Instrument Method:* | **IM_General0%** | *Bandwidth:* | **4** |
| *Processing Method:* | **Q10** | *Dilution Factor:* | **1.0000** |
| *Injection Date/Time:* | **25.Feb.21 12:52** | *Sample Weight:* | **1.0000** |

#### Chromatogram

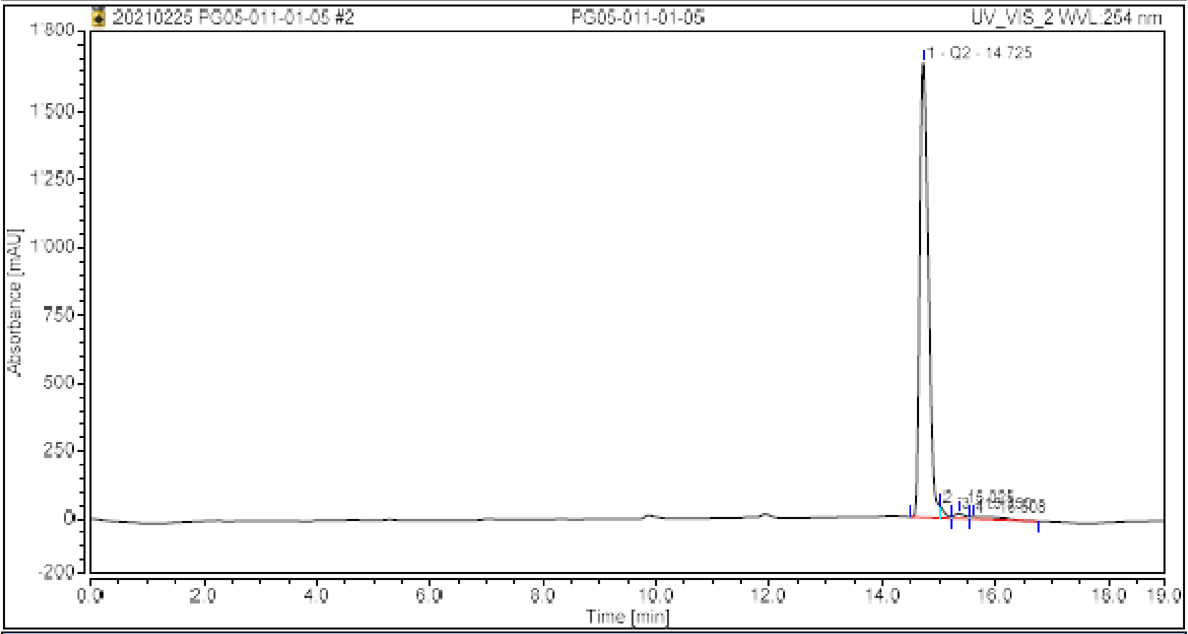

#### Integration Results

| No. | Peak Name | Retention Time min | Area mAU*min | Height mAU | Relative Area % | Relative Height % | Amount |
|---|---|---|---|---|---|---|---|
| 1 | Q2 | 14.725 | 291.487 | 1676.652 | 95.03 | 96.08 | n.a. |
| 2 | | 15.025 | 4.634 | 41.244 | 1.51 | 2.36 | n.a. |
| 3 | | 15.350 | 4.089 | 18.398 | 1.33 | 1.05 | n.a. |
| 4 | | 15.608 | 6.533 | 8.683 | 2.13 | 0.50 | n.a. |
| **Total:** | | | **306.743** | **1744.978** | **100.00** | **100.00** | |

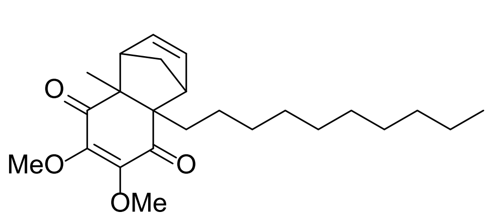

4a-decyl-6,7-dimethoxy-8a-methyl-1,4,4a,8a-tetrahydro-1,4-methanonaphthalene-5,8-dione
Chemical Formula: $C_{24}H_{36}O_4$

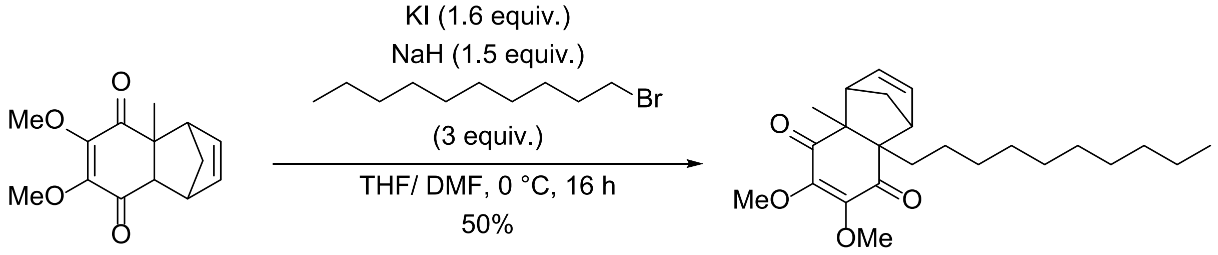

To a stirred solution of 6,7-dimethoxy-4a-methyl-1,4,4a,8a-tetrahydro-1,4-methanonaphthalene-5,8-dione (1.17549 g, 4.7346 mmol) in dry THF (2.5 mL) at 0 °C NaH (60% dispersion in parrafinum, 0.250 g, 6.4277 mmol) was added. The mixture was stirred for 45 minutes at 0 °C, giving a brown slurry. To this was added 1-bromodecane (2 mL, 2.132 g, 9.6391 mmol) and a solution of KI (0.250 g, 1.5058 mmol) in dry DMF (2.5 mL). The mixture was stirred at 0 °C for 15 minutes, afterwhich it was slowly warmed to 21 °C. Then additional KI (0.86345 g, 5.2008 mmol) was added, and the mixture was stirred for another 15 h, resulting in a brownish semi-solid mixture. To this was added a mixture of water and ethyl acetate with little $Na_2S_2O_3$. The aqueous phase was extracted with ethyl acetate, the combined organic layers were washed with brine, dried over $MgSO_4$, filtered, and concentrated under reduced pressure. The mixture was taken up in dichloromethane and concentrated onto silica gel. The crude was purified by flash column chromatography (cyclohexane/ ethyl acetate gradient from 1:0 to 4:1). The title compound was obtained in 50 % yield (0.79448 g, 2.04455 mmol) as orange oil.

**$^1$H NMR** (300 MHz, $CDCl_3$) $\delta$ 6.08 – 6.03 (m, 2H), 3.93 (s, 3H), 3.90 (s, 3H), 3.10 – 3.08 (m, 1H), 3.00 – 2.98 (m, 1H), 1.94 – 1.87 (m, 1H), 1.77 – 1.65 (m, 2H), 1.47 (s, 3H), 1.27 – 1.23 (m, 17H), 0.90 – 0.85 (m, 3H). **$^{13}$C NMR** (75 MHz, $CDCl_3$) δ 199.01, 198.56, 150.55, 149.63, 138.32, 137.27, 60.39, 60.32, 59.64, 56.37, 54.42, 52.81, 43.48, 37.43, 32.02, 30.54, 29.73, 29.68, 29.49, 29.44, 26.44, 23.39, 22.81, 14.25. **HRMS** (ESI) calculated for $[M+H]^+$ 389.2686, found 389.2682.

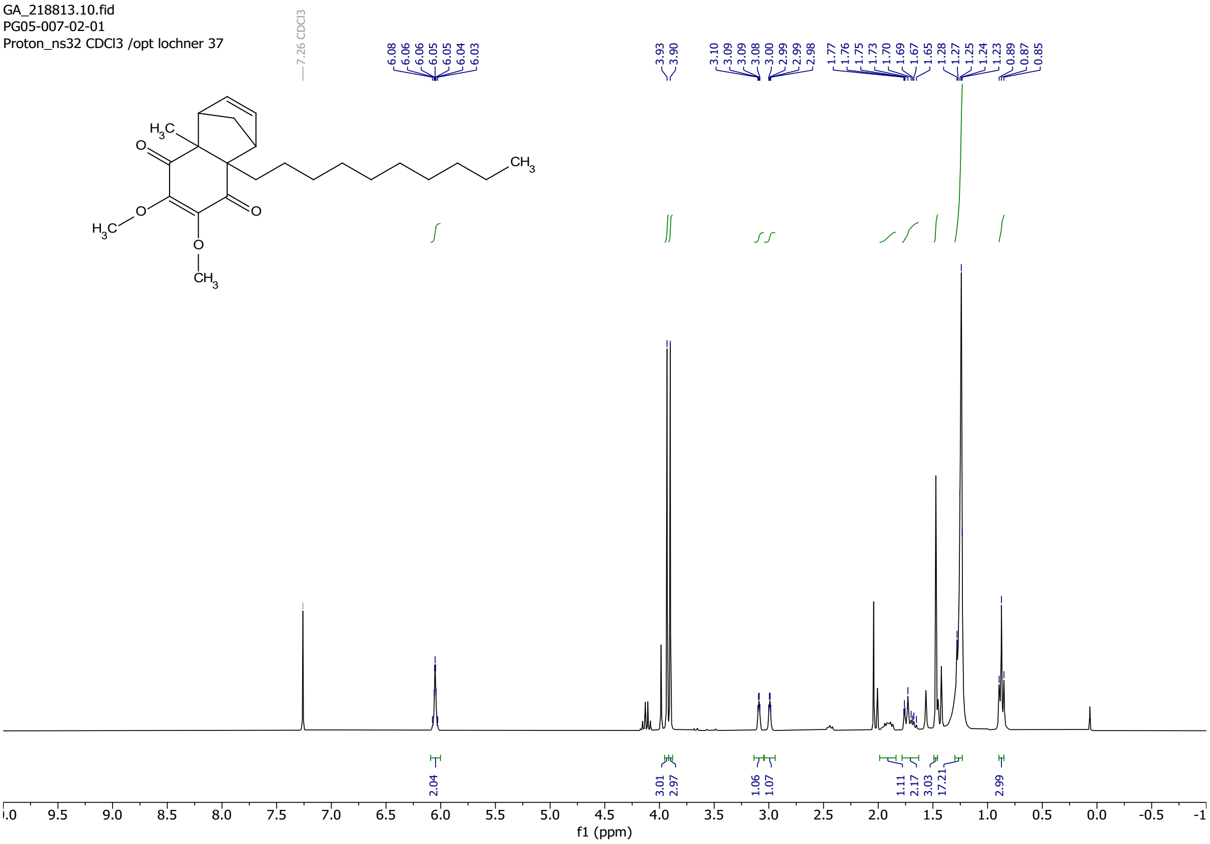
GA_218813.10.fid
PG05-007-02-01
Proton_ns32 CDCl3 /opt lochner 37
7.26 CDCl3
6.08 6.06 6.06 6.05 6.05 6.04 6.03
3.93 3.90
3.10 3.09 3.09 3.08 3.00 2.99 2.99 2.98
1.77 1.76 1.75 1.73 1.70 1.69 1.67 1.65 1.28 1.27 1.25 1.24 1.23 0.89 0.87 0.85
$H_3C$
O
$CH_3$
2.04
3.01 2.97
1.06 1.07
1.11 2.17 3.03 17.21
2.99
f1 (ppm)

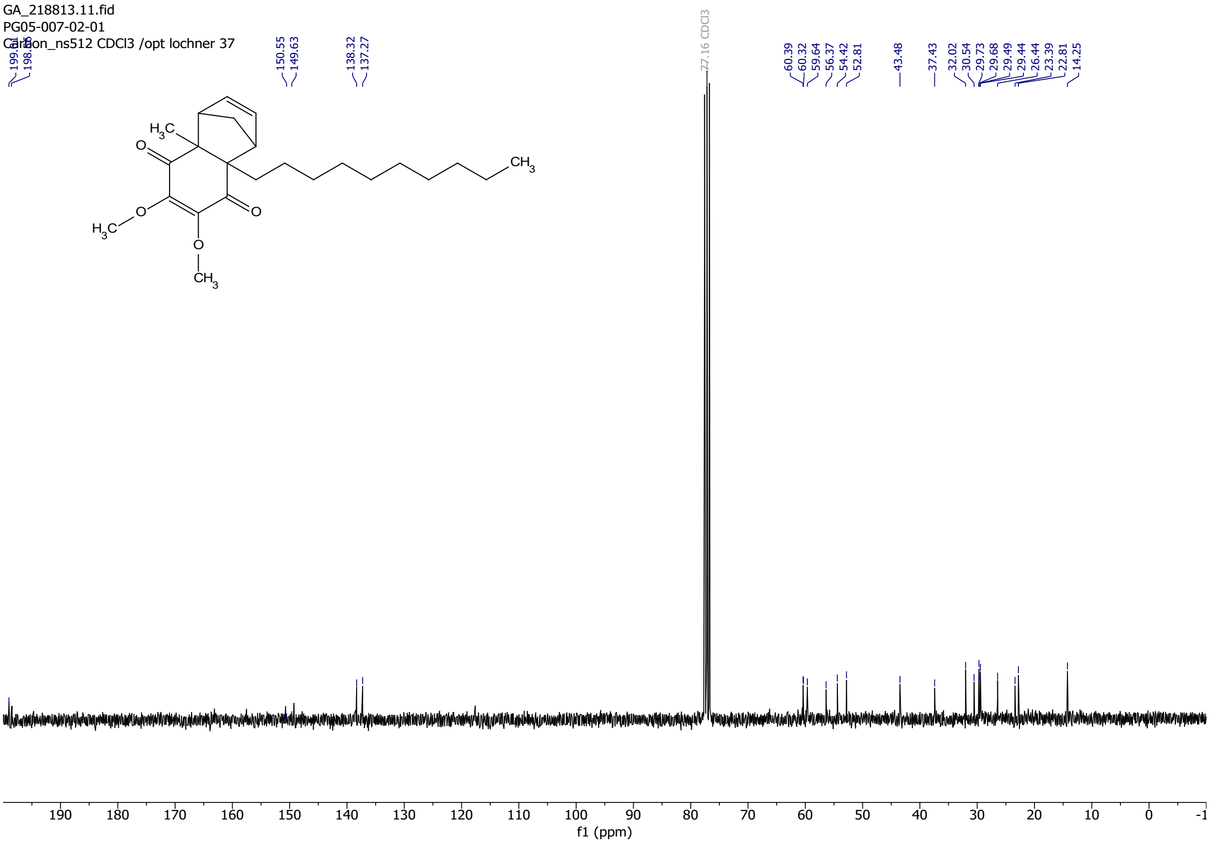
GA_218813.11.fid
PG05-007-02-01
Carbon_ns512 CDCl3 /opt lochner 37
150.55 149.63
138.32 137.27
77.16 CDCl3
60.39 60.32 59.64 56.37 54.42 52.81
43.48
37.43
32.02 30.54 29.73 29.68 29.49 29.44 26.44 23.39 22.81 14.25
$H_3C$
O
$CH_3$
f1 (ppm)

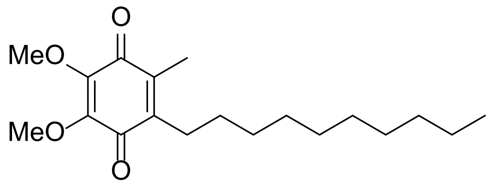

2-decyl-5,6-dimethoxy-3-methylcyclohexa-2,5-diene-1,4-dione
Chemical Formula: $C_{19}H_{30}O_4$

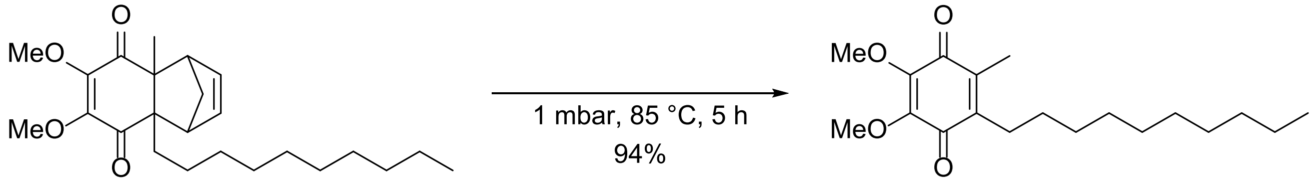

4a-decyl-6,7-dimethoxy-8a-methyl-1,4,4a,8a-tetrahydro-1,4-methanonaphthalene-5,8-dione (0.28314 g, 0.7287 mmol) was heated to 85 °C at ~1 mbar atmosphere for 5 h. The mixture gradually turned dark red. The red oil was diluted with dichloromethane and concentrated onto silica gel. The crude was purified by reverse phase flash column chromatography (C18, $H_20$ + 0.1 % TFA/ MeCN + 0.1 % TFA gradient from 1:0 to 0:1). The title compound was obtained in 94 % yield (0.62001 g, 1.9228 mmol) as red oil.

**$^1$H NMR** (400 MHz, $CDCl_3$) $\delta$ 3.98 (m, 6H), 2.44 (t, *J* = 7.4 Hz, 2H), 2.01 (s, 3H), 1.39 – 1.25 (m, 16H), 0.87 (t, *J* = 6.8 Hz, 3H). **$^{13}$C NMR** (101 MHz, $CDCl_3$) $\delta$ 184.87, 184.30, 144.50, 144.48, 143.28, 138.80, 61.27 (2C), 32.03, 29.98, 29.72, 29.66, 29.52, 29.44, 28.89, 26.55, 22.81, 14.24, 12.03. **HRMS** (ESI) calculated for $[M+H]^+$ 323.2217, found 323.2211.

**Purity (HPLC)**: 99 % (at 214 nm), 98 % (at 254 nm).

004382.10.fid
H1_STD CDCl3 /opt service 46

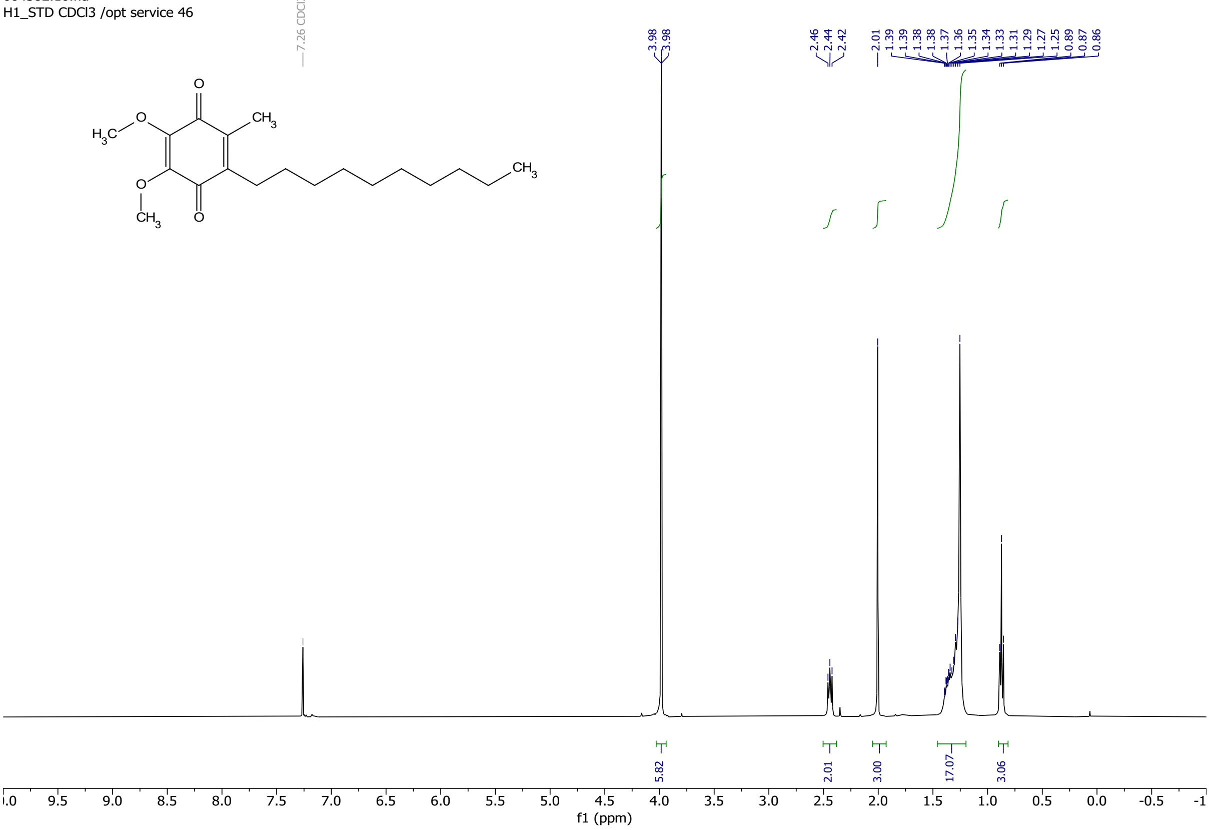

004382.11.fid
C13CPD_STD CDCl3 /opt service 46

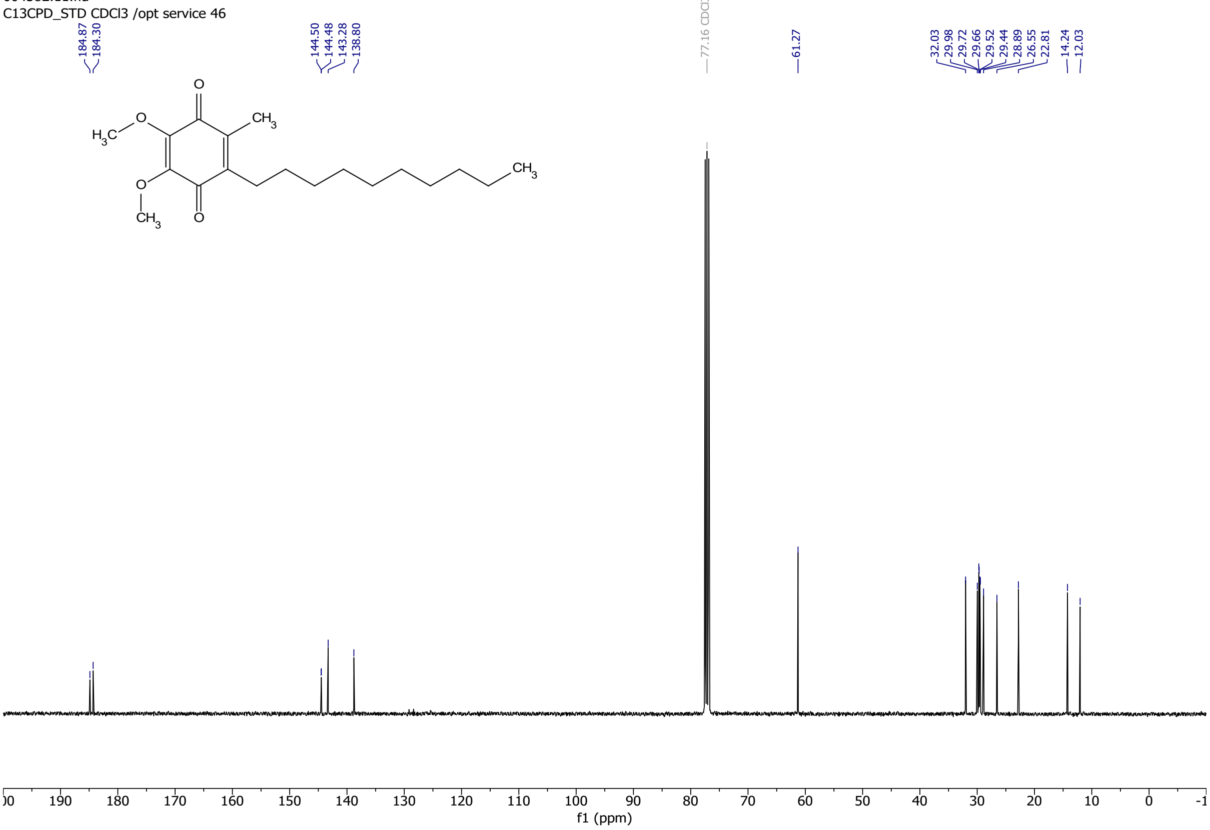

### Chromatogram and Results

#### Injection Details

| | | | |
|---|---|---|---|
| *Injection Name:* | **PG05-008-02-03** | *Run Time (min):* | **19.00** |
| *Vial Number:* | **BC3** | *Injection Volume:* | **5.00** |
| *Injection Type:* | **Unknown** | *Channel:* | **UV_VIS_1** |
| *Calibration Level:* | | *Wavelength:* | **214** |
| *Instrument Method:* | **IM_General40%** | *Bandwidth:* | **4** |
| *Processing Method:* | **Decyl** | *Dilution Factor:* | **1.0000** |
| *Injection Date/Time:* | **12.Mär.21 14:25** | *Sample Weight:* | **1.0000** |

#### Chromatogram

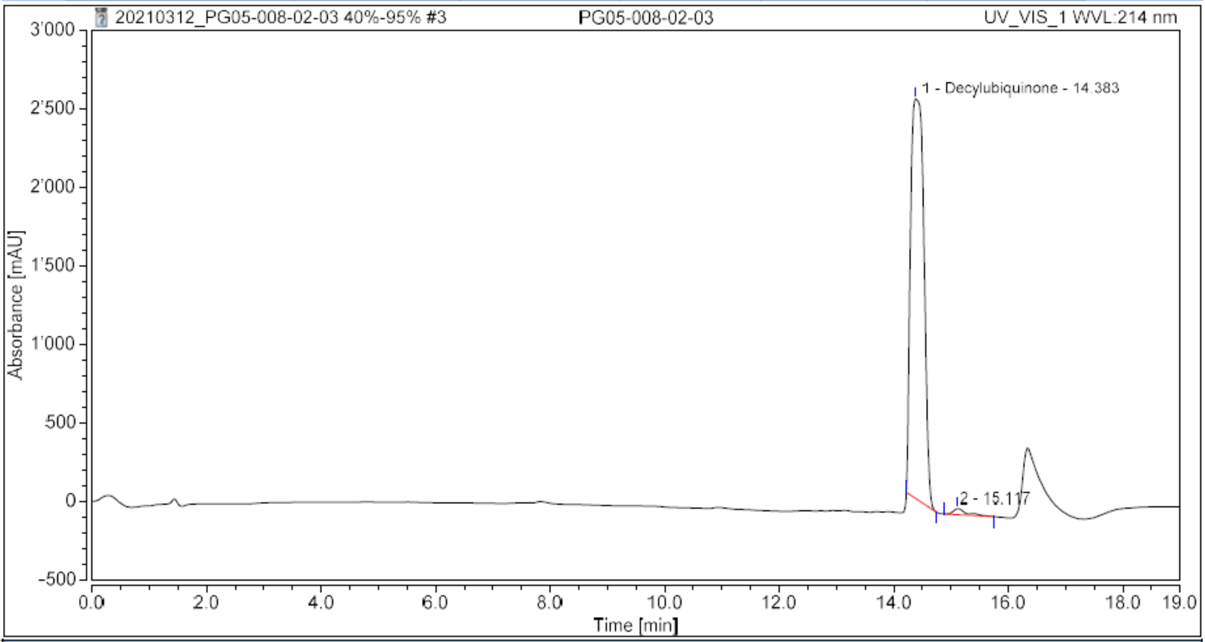

#### Integration Results

| No. | Peak Name | Retention Time min | Area mAU*min | Height mAU | Relative Area % | Relative Height % | Amount |
|---|---|---|---|---|---|---|---|
| 1 | Decylubiquinone | 14.383 | 687.345 | 2550.073 | 98.58 | 98.53 | n.a. |
| 2 | | 15.117 | 9.929 | 38.159 | 1.42 | 1.47 | n.a. |
| **Total:** | | | **697.274** | **2588.233** | **100.00** | **100.00** | |

#### Chromatogram and Results

**Injection Details**

| | | | |
|---|---|---|---|
| *Injection Name:* | **PG05-008-02-03** | *Run Time (min):* | **19.00** |
| *Vial Number:* | **BC3** | *Injection Volume:* | **5.00** |
| *Injection Type:* | **Unknown** | *Channel:* | **UV_VIS_2** |
| *Calibration Level:* | | *Wavelength:* | **254** |
| *Instrument Method:* | **IM_General40%** | *Bandwidth:* | **4** |
| *Processing Method:* | **Decyl** | *Dilution Factor:* | **1.0000** |
| *Injection Date/Time:* | **12.Mär.21 14:25** | *Sample Weight:* | **1.0000** |

**Chromatogram**

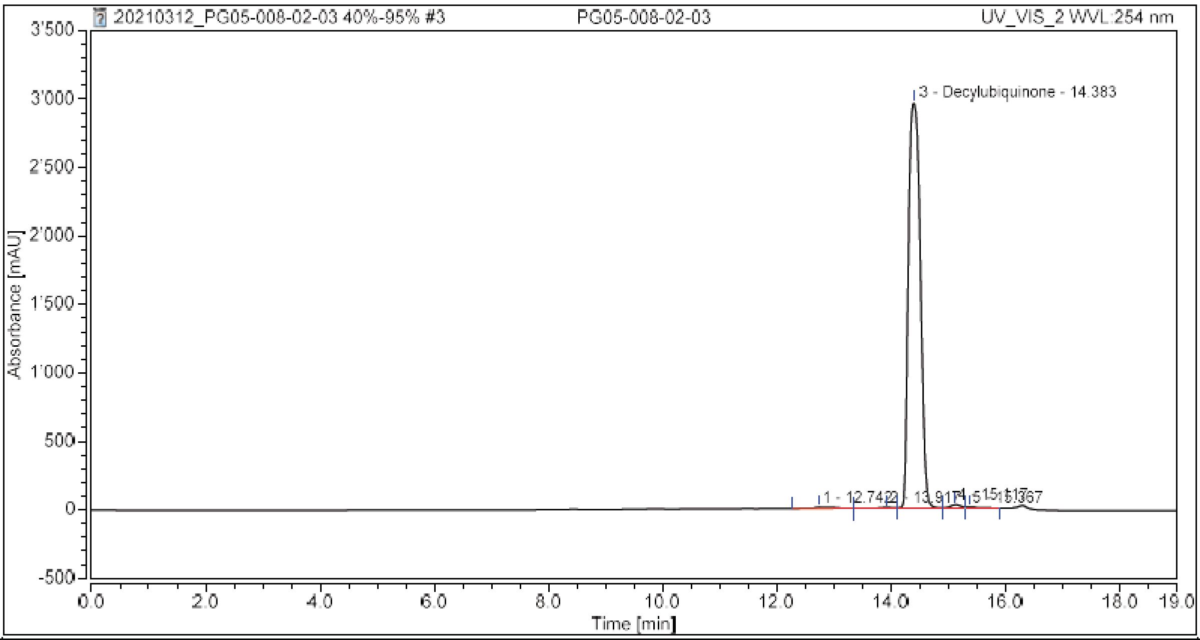

**Integration Results**

| No. | Peak Name | Retention Time min | Area mAU*min | Height mAU | Relative Area % | Relative Height % | Amount |
|---|---|---|---|---|---|---|---|
| 1 | | 12.742 | 3.638 | 8.099 | 0.50 | 0.27 | n.a. |
| 2 | | 13.917 | 2.057 | 6.593 | 0.28 | 0.22 | n.a. |
| 3 | Decylubiquinone | 14.383 | 711.205 | 2961.692 | 98.20 | 98.37 | n.a. |
| 4 | | 15.117 | 5.131 | 24.882 | 0.71 | 0.83 | n.a. |
| 5 | | 15.367 | 2.209 | 9.398 | 0.31 | 0.31 | n.a. |
| **Total:** | | | **724.241** | **3010.664** | **100.00** | **100.00** | |

1 Lu, L. & Chen, F. A Novel and Convenient Synthesis of Coenzyme Q1. *Synthetic Communications* **34**, 4049-4053, doi:10.1081/SCC-200036578 (2004).
